# Cell-free prototyping of AND-logic gates based on heterogeneous RNA activators

**DOI:** 10.1101/661561

**Authors:** François-Xavier Lehr, Maleen Hanst, Marc Vogel, Jennifer Kremer, H. Ulrich Göringer, Beatrix Suess, Heinz Koeppl

## Abstract

RNA-based devices controlling gene expression bear great promise for synthetic biology, as they offer many advantages like short response times and light metabolic burden compared to protein-circuits. However, little work has been done regarding their integration to multi-level regulated circuits. In this work, we combined a variety of small transcriptional activator RNAs (STARs) and toehold switches to build highly effective AND-gates. To characterise the components and their dynamic range, we used an *Escherichia coli* (*E. coli*) cell-free transcription-translation (TX-TL) system dispensed via nanoliter droplets. We analysed a prototype gate *in vitro* as well as *in silico*, employing parameterised ordinary differential equations (ODEs), where parameters were inferred via parallel tempering, a Markov chain Monte Carlo (MCMC) method. Based on this analysis, we created nine additional AND-gates and tested them *in vitro*. The functionality of the gates was found to be highly dependent on the concentration of the activating RNA for either the STAR or the toehold switch. All gates were successfully implemented *in vivo*, offering a dynamic range comparable to the level of protein circuits. This study shows the potential of a rapid prototyping approach for RNA circuit design, using cell-free systems in combination with a model prediction.

**Abbreviations:** TX-TL (transcription-translation), ODEs (ordinary differential equations), STARs (small transcriptional activator RNAs), MCMC (Markov chain Monte Carlo).

## Introduction

Over the last decade, RNA molecules have emerged as one of the most promising elements in the toolbox of synthetic biology. They are able to modulate almost all cellular processes, including transcription, translation, and mRNA degradation or splicing (*1*, *2*). This in turn has led to the engineering of new regulatory RNAs, either derived from natural molecules or based on novel mechanisms, with a great diversity of functions and targets. These regulators have been successfully used to address current problems in biotechnology, such as challenges in metabolic engineering (*3*, *4*) or therapeutic advances (*5*, *6*). Regulatory RNAs possess many features making them particularly suitable for genetic circuit design with advantages over transcription factors. One advantage is that RNA is able to propagate information solely at the transcriptional level, without the need for intermediate proteins, which reduces the metabolic burden of a cell (*7*) and increases signal propagation speed (*8*). Different RNA regulatory devices like riboswitches (*9*), ribozymes (*10*), small transcriptional activators (STARs) (*11*), toehold switches (*12*) as well as CRISPR/Cas based systems are now available for building synthetic logic circuits. Lee et al. have recently constructed logic gates in *E. coli* based on the combination of CRISPR/antisense RNA (asRNA) and STARs/asRNA (*13*, *14*). Shen et al. have built layered AND-gates out of engineered sensor RNA, offering modularity at the post-transcriptional level (*10*). These examples show the great potential of RNA devices, but also highlight some of the challenges in assembling highly complex synthetic circuits. The dynamic range, for instance, is often a limiting factor when tight regulation is needed. Another problem is that advanced applications require a variety of well characterised and orthogonal devices. One way to address these challenges is to use RNA-based circuits which are able to regulate gene expression at different levels. However, little work has been done yet with respect to the integration of different RNA regulators in the same circuit (*15*, *16*).

In this work, we built and tested synthetic gene constructs that were regulated by both STARs and toehold switches. Similar to protein-based systems, STARs and toehold switches regulate gene expression with very high dynamic ranges and great specificity for their targets, and are available in libraries of orthogonal parts or design rules for creating new ones (*12*, *17*). Additional reasons led us to select STARs and toehold switches. First, they are able to control transcription and translation independently, offering solutions for multiplexed gene regulation. Secondly, using such a multi-level regulator system has been shown to increase the dynamic range of gene expression (*18*). Thirdly, both systems have been thoroughly characterised with fine-tuned sequences or coupled with various synthetic parts, turning them into candidates of choice for integrated circuits (*19*, *20*). Finally, because both switching mechanisms are based on Watson-Crick base pairing, gate design could be aided by the growing number of algorithms for computing secondary structures (*21*, *22*).

In this study, we performed rapid prototyping by using an *E. coli* TX-TL system. Cell-free protein synthesis has emerged recently as an ideal platform for rapidly prototyping synthetic parts or novel circuits within hours (*23*, *24*) in a high-throughput manner (*25*). The use of a liquid-handling robot enables us to downscale dispensing to the nano-litre range, reducing cost and preparation time, as well as improving reliability of the resulting data (*26*).

We started with two individual systems: a transcriptional regulator based on a STAR (AD1) and a translational regulator based on a toehold (toehold 2). For each system, we set up an ODE-model and used Bayesian inference to obtain the model parameters. We merged the two models and their parameters to predict the dynamics of our prototype STAR AD1-toehold 2 gate. The merged gate-model indicated the need for tight transcriptional components and allowed an estimate of suitable concentrations for future gates.

Based on those *in silico* insights, we built five additional single activators and nine AND-gates and characterised them by titration experiments. All gates showed robust logical output for any combination of input. Finally, we implemented the constructs in *E. coli*, thus verifying the functionality of each gate *in vivo*. The results show that cell-free TX-TL systems are able to screen functional candidates and therefore can rapidly prototype novel RNA-based circuits. Our findings provide a basis for an RNA-circuitry framework in TX-TL systems, where optimisation of DNA part concentration is critical for achieving a maximum dynamic range in an environment where resources are limited.

## Results and discussion

### Characterisation of RNA activators

To build multilevel RNA-regulated AND-gates based on the STARs and toehold switches, our first step was to assess the functionality of the individual activator components in TX-TL. STARs have been designed based upon the conditional formation of RNA-terminator hairpins. Synthetic STAR antisense sequences are used to disrupt the formation of these intrinsic terminators placed in a target module upstream of the regulated gene. In the absence of the anti-terminator STAR, the STAR-target leads to an early rho-independent termination of the regulated transcript. The toehold system controls gene expression on the translational level and also contains an antisense RNA which is called trigger. In the absence of the trigger, the toehold blocks the ribosomal binding site (RBS), while binding of the trigger to the toehold leads to a structure where the RBS is accessible for the ribosome and translation can occur. Thus both devices become active in the presence of an interacting antisense RNA (Fig. 1 A).

**Figure 1:**
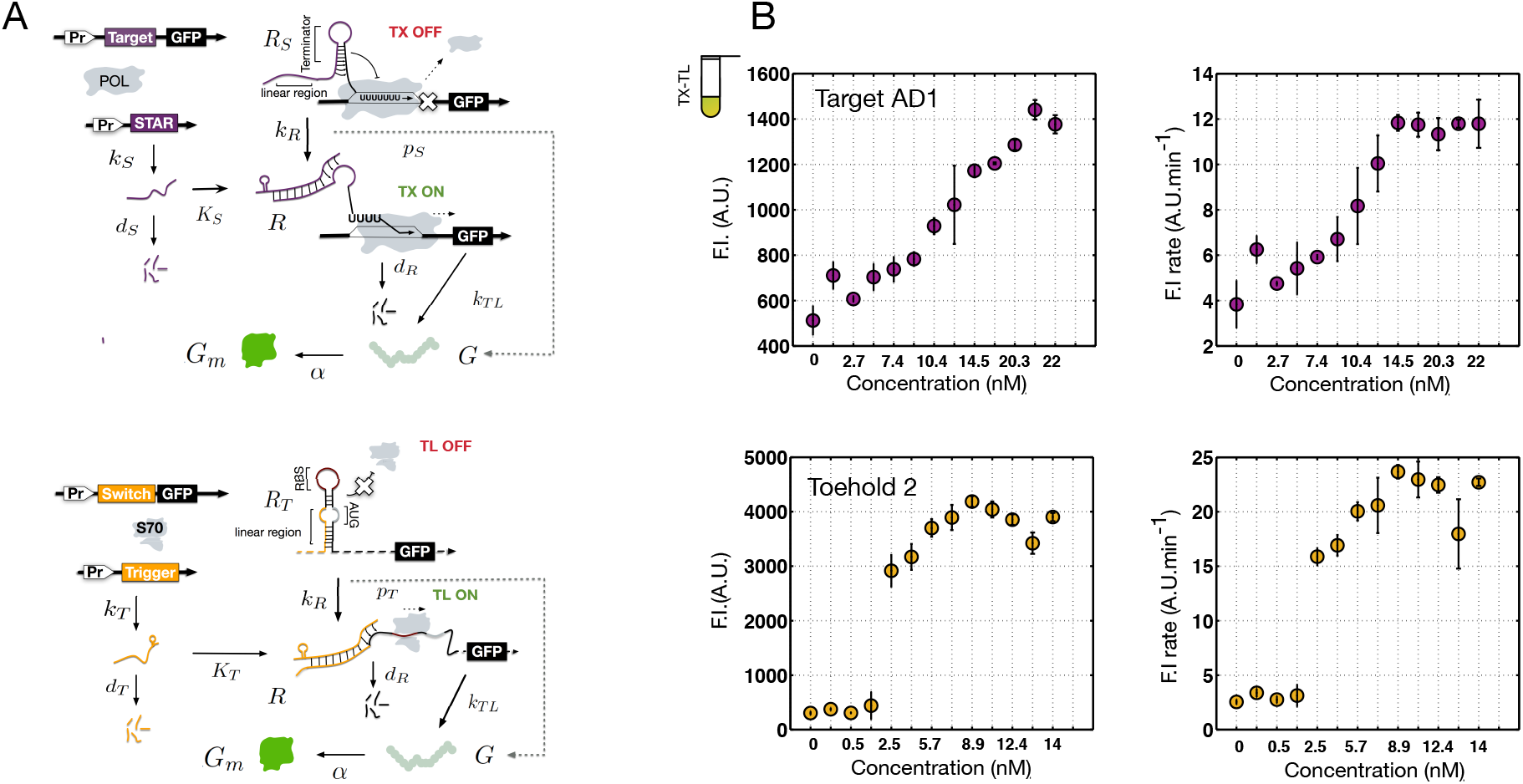
TX- and TL-regulation. (A) Mechanism of the small transcriptional activator RNAs regulating the transcriptional activity of the targeted gene; mechanism of the RNA-based toehold switch regulating the translation of the gene. (B) Corresponding dose response curves of TX-TL reactions with 2 nM of regulated reporter plasmid and titration of DNA-encoded trigger or STAR. Endpoint fluorescence measurements (left) and sfGFP production rates (right) are presented. Rates were determined by computing the slope in the linear regime of each time-course trace. Errors bars represent the standard deviation over three independent experiments.

We chose the same two plasmid system as used by Chappell et al. (*11*). One plasmid contained the activating RNA, which is driven by the strong constitutive J23119 promoter. The other plasmid codes for the super folder GFP (sfGFP) (*27*) reporter gene, which is also driven by the J23119 promoter. Either the STAR target or the toehold switch sequence is placed upstream of the reporter gene to control its expression. In both plasmids, the t500 terminator is used to terminate transcription. For the *in vivo* experiments, the activator cassettes were combined in the same plasmid.

We characterised the expression of the most efficient STAR from the original publication (STAR AD1) (*11*) together with one of the top-performing engineered toehold switches (toehold 2) (*12*). Dose response curves were generated for each part using the TX-TL system by varying the concentration of the activator plasmid. We used a volume of 2 μl by dispensing cell extract, buffer and plasmid DNA with the help of a liquid handling robot. The reporter plasmid concentration was 2 nM, and we varied the concentration of activator plasmid (trigger 2 or STAR AD1) between 0-22 nM. The results of the endpoint measurements are presented in Fig. 1 B and individual trace kinetics are available in Suppl. Fig. 5.

The TX-regulation-system and the TL-regulation-system reached saturation at about 20 nM of STAR plasmid and 7 nM of trigger plasmid, respectively. Similar to previous *in vivo* data (*11*, *12*), the toehold 2 has a three times higher dynamic range than the STAR AD1. Rather than due to impaired functionality, those lower dynamic ranges could be attributed to the TX-TL system. Previous results have highlighted the decrease in dynamic ranges for RNA devices when transitioning from *in vivo* to *in vitro* (*28*). The dose-response relationships follow either a linear response for STAR AD1 or a fast saturation curve for toehold 2 activation. This difference may be explained by the longer time in which a toehold strand can be displaced by a trigger RNA (mRNA life-time) compared to the transient event characterising the STAR termination (*19*). To optimise the sfGFP read-out (Suppl. Fig. 6), we increased the reporter plasmid concentration to 6 nM. This led to a significant increase in translated sfGFP. However, the concentration of activator plasmid needed to reach the plateau tripled from 7 nM to 20 nM, suggesting that RNA polymerases were not saturated yet with 2 nM of DNA template. Because such high DNA concentrations caused issues with dispensing, the reporter plasmid concentration was kept at 2 nM. The dynamic range of around 20-fold was unaffected by the different concentration levels.

### Mathematical modeling

Based on the data for the STAR AD1 and the toehold 2 switch from the TX-TL experiments, we built coarsegrained ODE-models for both individual systems. The eventual goal was to use the parameters from those individual models to predict the performance of the AND-gate.

For each model, we restricted the number of parameters and species to account for the unidentifiability of the parameters because only one species was measured. The models were mainly based on mass-action kinetics except for the TX- and TL-regulation, for which we employed Hill-kinetics to capture the corresponding dose-response shapes. The detailed models can be found in the Supplement.

Having set up the models, we calibrated the kinetic model curves of the mature sfGFP of the two single parts to the experimental data. We achieved this through a Bayesian inference method with Markov chain Monte Carlo sampling. To improve the convergence speed compared to standard approaches, we performed parallel tempering, a modification of the Metropolis-Hastings algorithm (for details see Sec. 3 Material and Methods). Each part was calibrated to the experimental data corresponding to different input concentrations of activator plasmid. The resulting parameter posterior distributions for the toehold-trigger 2 system are shown in Fig. 2 and the corresponding model kinetics are shown in Fig. 3. The results for the target-STAR AD1 system can be found in the Supplement, Fig. 4 and Fig. 3.

**Figure 2:**
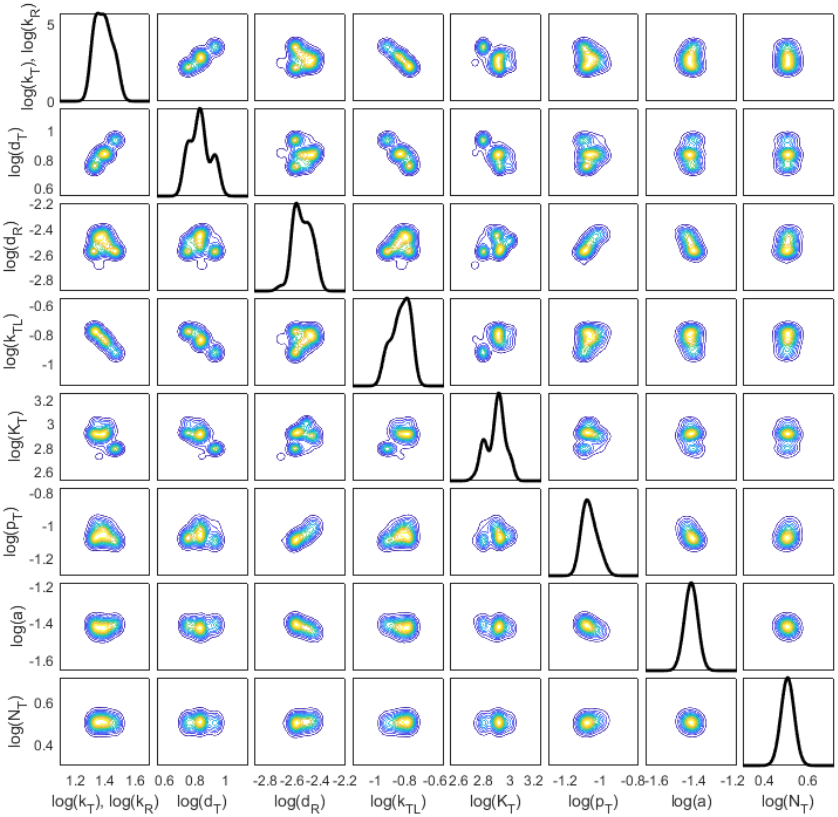
Parameter posterior distributions of the toehold-trigger 2 system based on 150 000 samples in the logarithmic space. Univariate distributions are on the diagonal, bivariate distributions on the off-diagonal. The blue colour at the boundary corresponds to low density values, the yellow colour to high values (relative to each other). Trace plots of the parameters were used to determine the burn-in phase and to ensure that convergence was reached. Further, a kernel density estimate with a Gaussian kernel was used.

**Figure 3:**
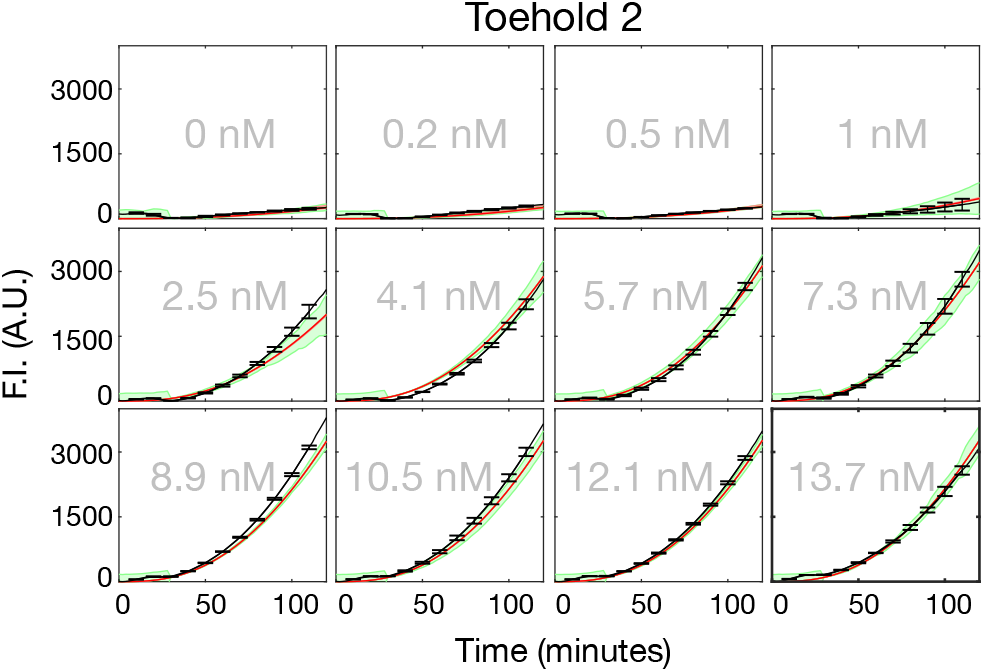
Calibration of single part: posterior predictive distribution of the toehold-trigger 2 system for 2 nM input concentration of toehold-sfGFP plasmid, corresponding to the posteriors in Fig. 2. Gray in the background: input concentrations of trigger plasmid; black: experimental data together with error bars at selected time points; red: median of parameterized model according to identified parameter posterior distribution; green: observation model corresponding to experimentally measured standard deviation and assumption of normally distributed observation errors, 95%-quantil and 5%-quantil.

**Figure 4:**
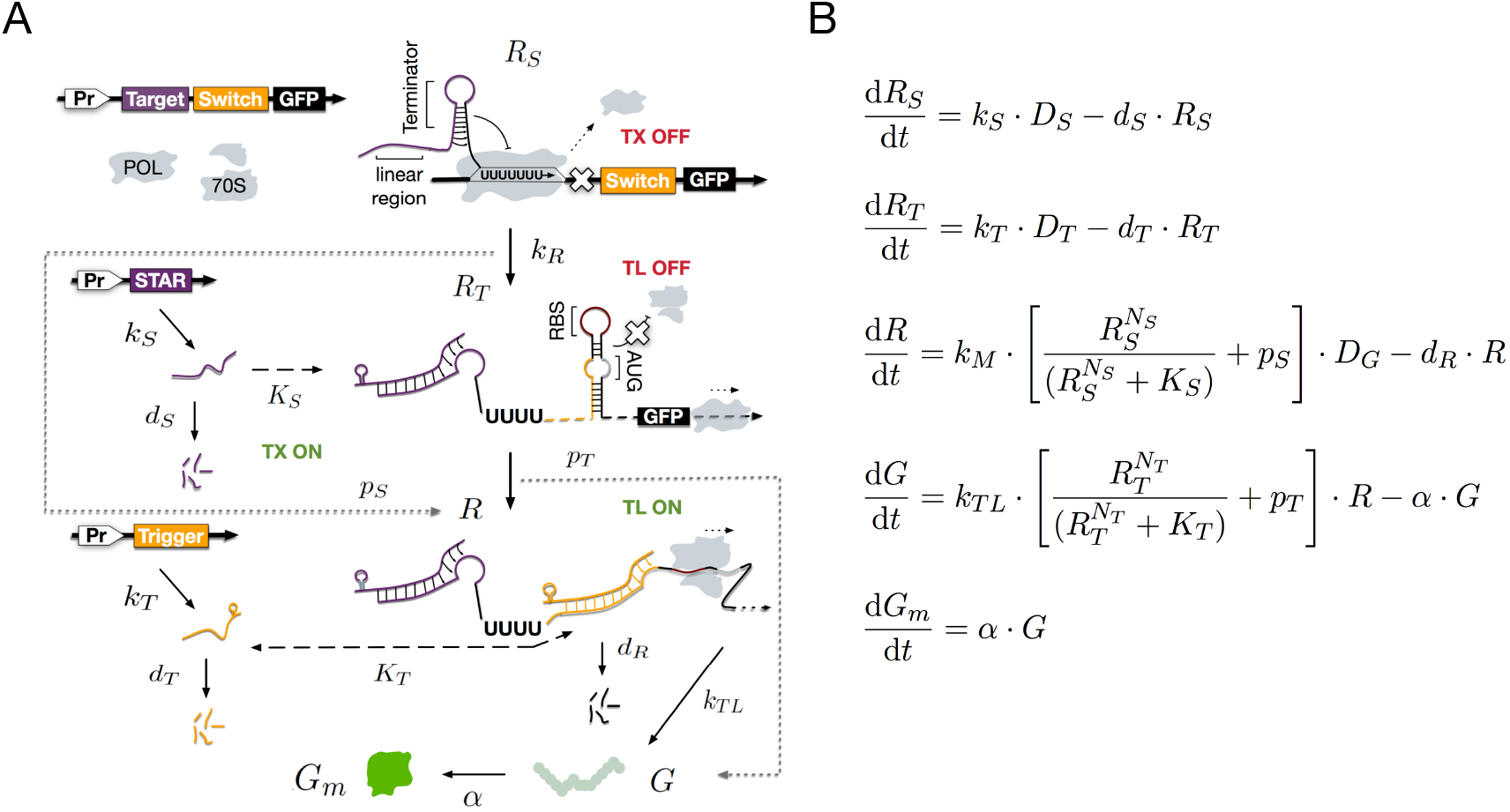
Scheme of the computational AND logic based on transcriptional (TX) and translational (TL) control. TX is regulated through the presence or absence of a STAR activator.TL is regulated through the binding of the trigger RNA to the linear toehold region. Both types of small RNA are required to activate the gene expression. Parameterized ODE-model of the AND-gate as a combination of the single part models. Besides standard mass-action kinetics, the model includes Hill kinetics for the TX- and TL-regulation mechanism. Details regarding the parameters, species and single part models can be found in the Supplement.

From the pairwise joint posteriors of the toehold 2 system (Fig. 2) we observe a positive correlation between the transcription and degradation rate of the trigger mRNA, while the transcription and translation rate of the reporter are negatively correlated. This matches expectations, since the higher the TX-rate of the reporter mRNA is, the lower its TL-rate should be to reach the measured fluorescence.

Our model is able to capture the steep increase between 1 nM and 4.1 nM input concentration of activator plasmid (see Fig. 3). This is possible due to the Hill-kinetics we used to model the regulation (c.f. Supplement).

Finally, we built the AND-gate model (Fig. 4) by combining the single part models and their inferred parameters (maximum a posteriori values). The model prediction of the gate can be seen in Fig. 5. In particular, we found that the model predicts a high leakage for 0 nM STAR AD1 plasmid input concentration.

**Figure 5:**
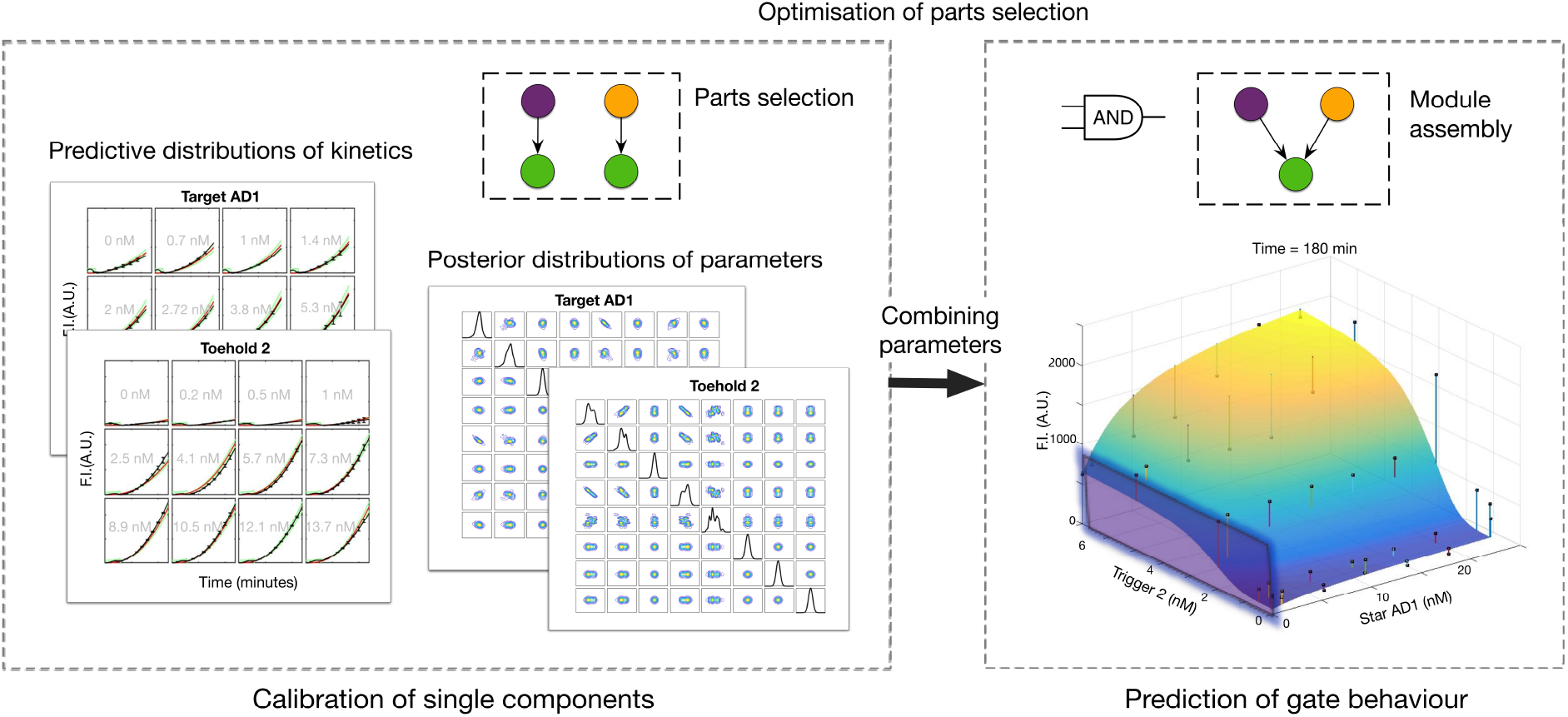
Computational dose-response of AND-gate (right) based on the combination of the parameters identified by the calibration of the single parts; AD1 target and toehold 2 (left). Qualitatively, the gate model predicts high leakage for the AD1 activator plasmid concentration set to 0 nM. Quantitatively, at least a rough prediction could be verified by the experimental data (black dots).

### Experimental evaluation of prototype gate-model prediction

After analyzing the AD1 target-toehold 2 gate *in silico*, we verified the predictions in TX-TL. The STAR AD1 target and the toehold 2 switch were placed in tandem upstream of a sfGFP reporter gene without further re-engineering (See Table 1). Based on the model, a combination of 42 concentrations was tested (six concentrations for the trigger and seven for the STAR). The predictions of the AND-gate model capture the qualitative trend of the experimental characterisation (Fig. 5). The TX-TL end-points fit to the predicted values of the sfGFP leakage along the STAR concentration gradient when no trigger is present. Some qualitative disagreements are observed at intermediate concentrations of activators, i.e. on the surface slope.

Finally, we wanted to assess if the predicted leakage *in vitro* could also provide insights into the *in vivo* behavior. Therefore, *E. coli* cells were transformed with the same reporter plasmid as used for the TX-TL measurements. This plasmid contains the p15A origin of replication, which leads to a low plasmid concentration inside the cell. For the activators, a plasmid with the ColE1 origin of replication was used, which leads to a high copy number inside the cell. To analyse the behaviour of the AND-gate, the *E. coli* cells were cotransformed with an activator plasmid containing either the STAR, the trigger or both activators on the same plasmid. The obtained endpoint fluorescence measurements show a similar behaviour of the AND-gate *in vivo* and *in vitro* (Supp. Fig. 1 A). We observe a slight activation of the gate when only trigger 2 is present, i.e., the transcriptional leakage observed *in silico* and *in vitro* occurred as well in living cells. This indicates that the use of TX-TL as a prototyping platform is not only beneficial for screening various circuit implementations, but can also be used to analyse more finely the effects of single part behaviour.

### Additional parts characterisation

As our prototype gate showed potential for proper functionality in TX-TL and *in vivo*, we sought next to optimise it by screening combinations of parts. Multiple studies have shown that combining single components may result in co-folding or context effects, challenging the implementation (*29*, *30*). However, STAR and toehold components are available as orthogonal matrices of numerous parts, and are constantly under improvement (*19*, *31*). Since we identified the relatively high OFF-stage of the STAR AD1 component as the source of translational leakage, we selected three newly engineered STARs with lowered OFF-levels (STAR 5, 6, STAR 8). In addition, we selected two other high performing toehold switches (toehold 1 and toehold 3) in order to increase the diversity of our gate library.

The five new RNA activators were tested under the same conditions as the initial parts in TX-TL. The corresponding dose-response curves of the sfGFP production rates are plotted in Fig. 6 A and individual traces are available in Supp. Fig. 5. The data also show that less trigger plasmid is needed to activate the toehold than STAR plasmid is needed to activate the STAR target. This was already observed for the STAR AD1 and toehold 2. No additional increase of the sfFGP production rate is seen after around 3 nM to 5 nM of trigger-encoding plasmid and after around 15 nM to 20 nM of STAR-encoding plasmid. Moreover, the OFF-level in the absence of the activator STAR DNA (5,6,8) was lower than the one from STAR AD1. The dynamic range increased as well, in particular for the STAR 6 construct. To build different AND-gates with the single parts, we verified *in vivo* whether the STAR 6 activates the toehold or if the trigger is able to activate the STAR target. This was not the case for any of the tested combinations (Supp. Fig. 2).

**Figure 6:**
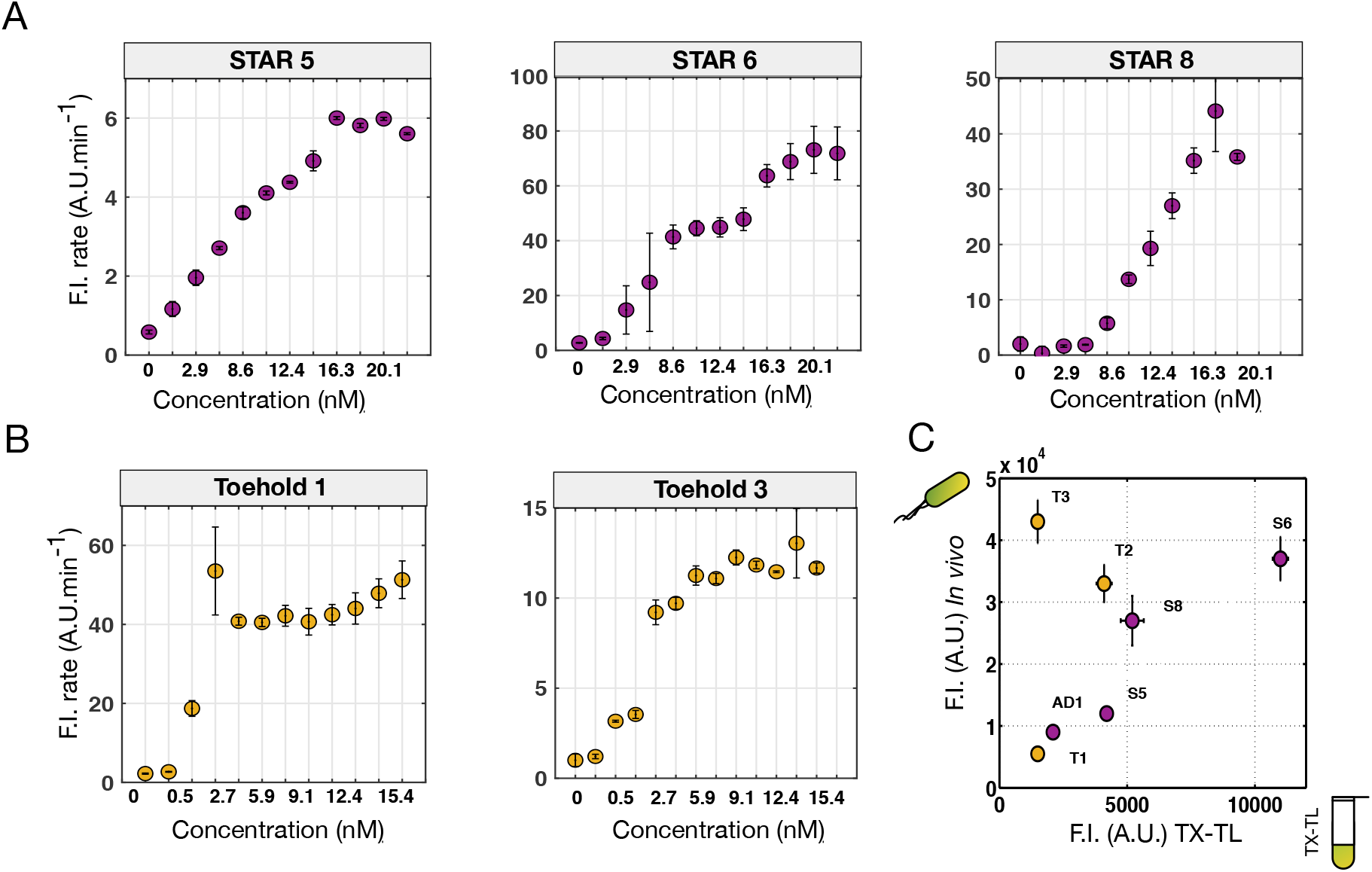
Additional STARs and toehold switches characterisation. (A) Dose-response curves of sfGFP production rates for STARs 5, 6, 8 in TX-TL. (B) Dose-response curves of sfGFP production rates for toehold 1 and 3. DNA-encoding STAR or trigger activators are titrated in presence of 2 nM of regulated sfGFP plasmid. Rates were obtained by calculating the slope in the linear regime of each time-course trace. (C) Corresponding relative fluorescence intensities between *in vivo* and *in vitro* for seven single parts. Errors bars represent the standard deviation of three independent experiments, for *in vitro* and *in vivo* data.

### Cross-design of single components into logic devices

We tested nine gates based on the cross design of the previous single components. This was done in two stages. First, a screening to detect the gates with the best dynamic ranges was performed. The concentrations were chosen according to the AD1-toehold 2 gate characterisation to pick up an active state (15 nM of STAR, 5 nM of trigger). Second, we performed a detailed analysis of the selected gate. Fig. 7 A displays the resulting kinetics over a period of 5 hours of expression. For every gate, the plateau is reached between 3 and 4 hours. All gates show functionality, ranging from around 4-fold for the least efficient (S5T2) to 74-fold for the most efficient (S6T3). Remarkably, S6T3 presents an absolute fluorescence intensity and a dynamic range almost one order of magnitude above the pool. Most of the gates display a symmetrical leakage when only trigger or STAR is expressed, except for S6T2, which shows higher leakage when only STAR is present.

**Figure 7:**
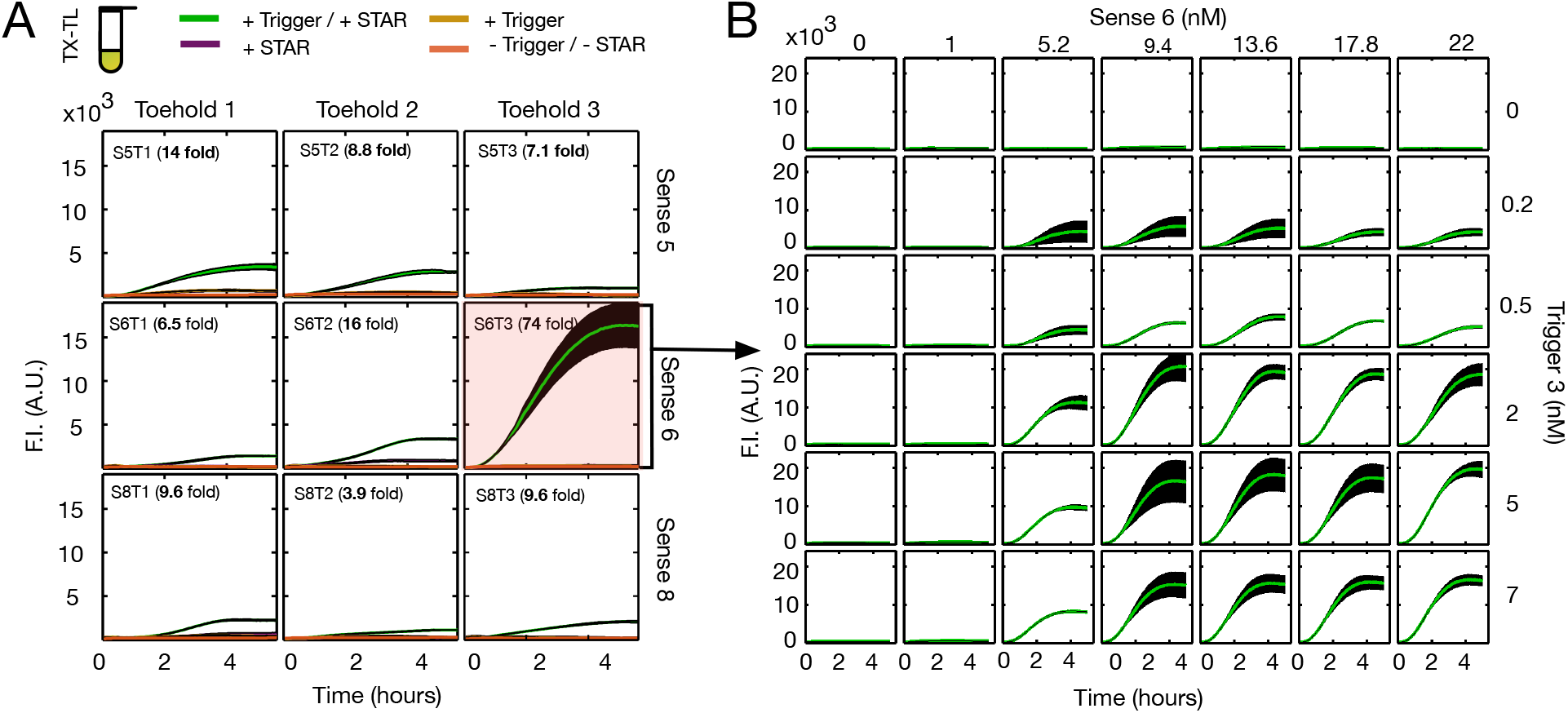
Characterisation of RNA-based logic AND-gates with TX-TL time course reactions. (A) Pre-screening in TX-TL reactions of RNA-based gates composed of a cross design of three toeholds (1, 2, 3) and three STARs (5, 6, 8), for nine gates. The black shaded regions correspond to the standard deviation over three independent reactions. Colored lines correspond to the average measurement over those replicates, according to the presence of either trigger-encoding DNA or STAR encoding DNA or both. The fold-range is computed as the difference between the active state (both trigger and STAR are present) divided by the inactive state (trigger and STAR are not present), corrected by the background value of the TX-TL mix. (B) Concentration matrix of TX-TL time course reactions for the sense 6-toehold 3-sfGFP gate (S6T3). Trigger and STAR encoding plasmids were simultaneously titrated from 0 nM to 7 nM and from 0 nM to 22 nM, respectively.

Correlation between gate performance and part composition has not been clearly identified. Our assumption was that for all constructs, the final sfGFP level could only be as high as the least efficient activators of the gate, i.e. the STAR or toehold. This can be observed for most of the gates, for instance, the gate S5T3 is one with the lowest fluorescence value – and is based on the two single components with the least efficient yield (target 5 and toehold 3, cf. Supp. Fig. 5). However, the S6T3 gate is a remarkable counter-example as the high yield of target 6 is not diminished by the presence of the toehold 3 component. We then performed kinetic measurements for S6T3 with similar conditions as the initial gate (Fig. 7 B). Initial screening concentrations were close to the optimal expression level, which were found to be 2 nM of trigger and around 9 nM of STAR.

### Comparison of dose-response matrices

To assess the qualitative disparity among the gates’ dose-responses, we performed kinetic measurements for two additional screened gates. The results are shown in Fig. 8. Compared to S6T3, the fluorescence levels were either intermediate (S6T2) or very low (S5T2). Overall, the qualitative shape remained similar. The initial screening concentrations were again near the maximum sfGFP-expression. A decrease in sfGFP fluorescence is observed when both activators are present in high concentration. This may be due to the toxicity effects reported for high DNA concentration in TX-TL (*32*). Resource competition is critical in TX-TL where the energy supply is not replenished. Monitoring transcription would be possible by tagging the sfGFP 3’ untranslated region (UTR) with a fluorescent reporter aptamer (*33*).

**Figure 8:**
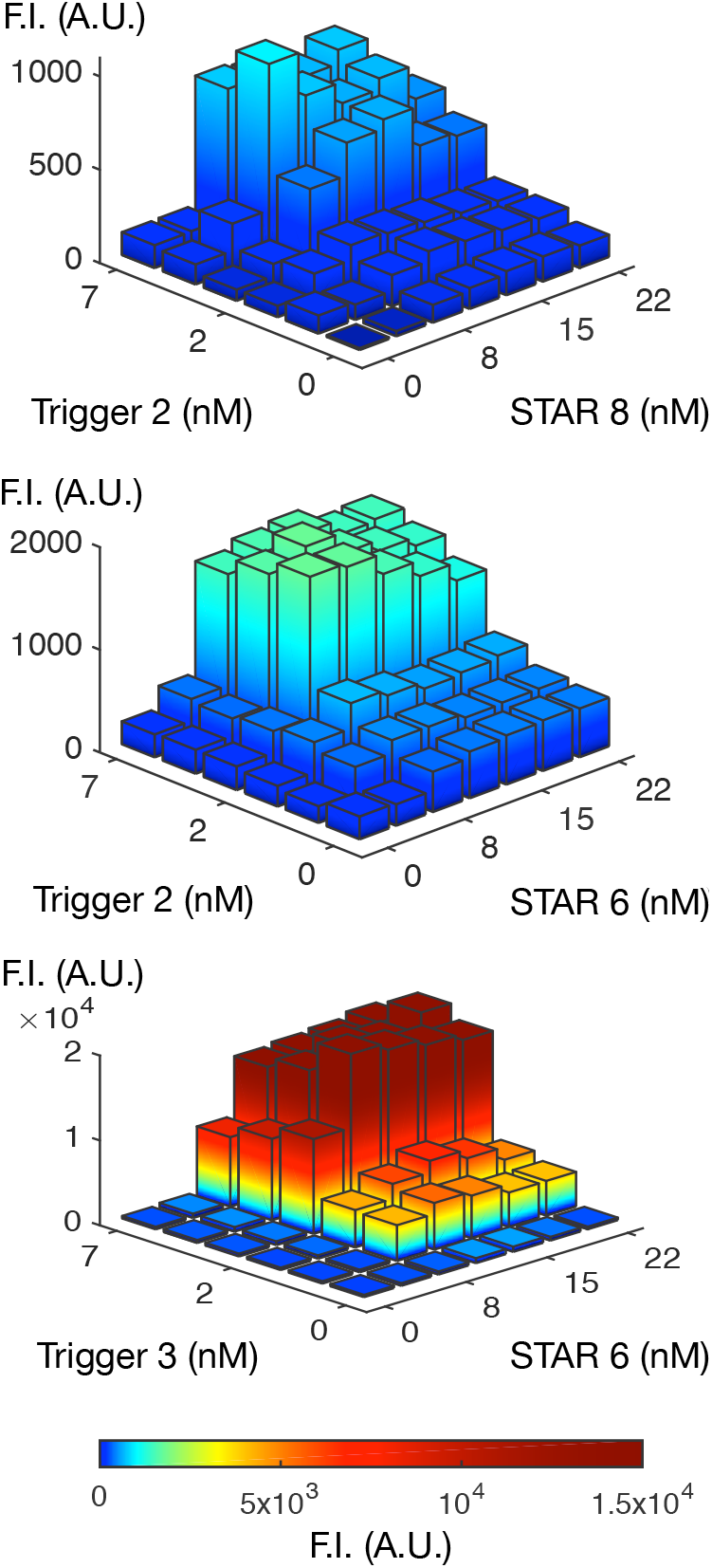
Dose-response comparison of endpoint fluorescence measurements of three different AND-gates after 4 hours of expression; sense 8-toehold 2, sense 6-toehold 2, and sense 6-toehold 3.

We investigated if *in silico* folding simulations could explain the difference of the sfGFP production between S6T3 and the other gates. Free energies of the RBS regions were computed for a range of sub-optimal structures (Supp. Fig. 9), showing comparable results for gates with the same toehold part. Minimum Free Energy (MFE) structures of the gates’ tails are displayed in Supp. Tab. 4. One can observe independent folding of the STAR and toehold secondary structures. The toehold component was not disrupted by the upstream spacer and STAR target terminator. We used the RBS-calculator tool (*34*) to analyse out-of-frame upstream start codons, but no difference between gates sharing the same toehold sequences was detected. We suspect that the exceptional S6T3 performance is due to the co-folding effect of the gate sequence on the sfGFP mRNA, which either affects the translation process or the mRNA stability (*35*).

**Figure 9:**
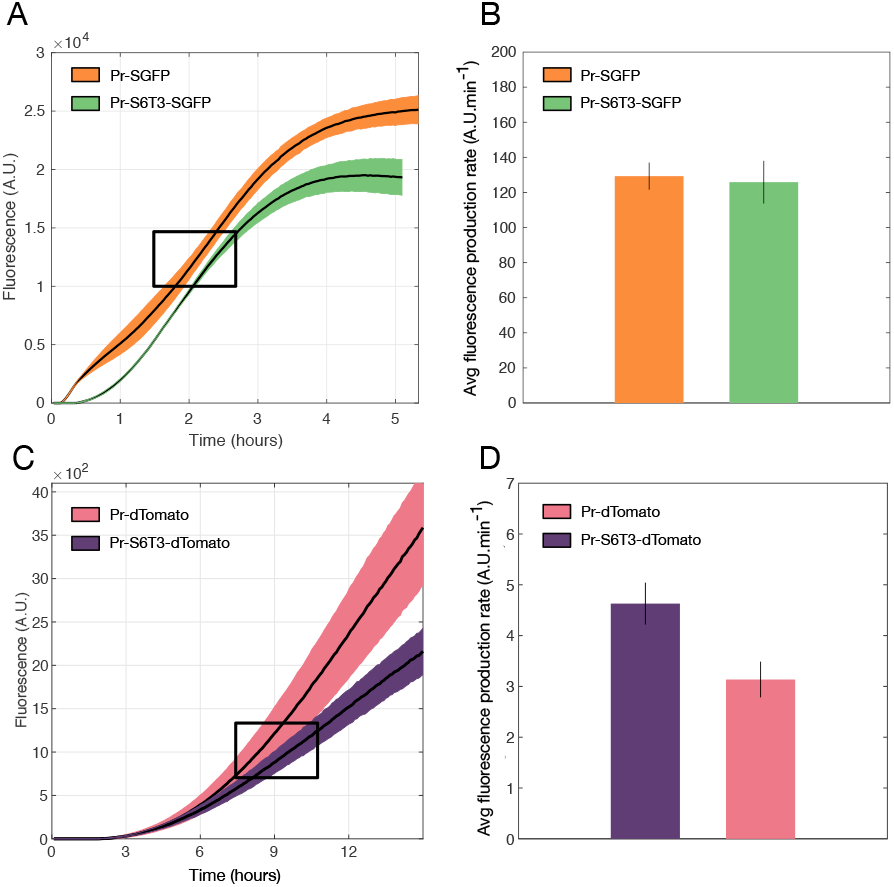
Comparisons between the S6T3 gate and unregulated fluorescent reporters. A, C: Fluorescence time courses of TX-TL reactions containing either 2 nM of DNA encoding sfGFP/dTomato with no RNA regulation, or with 2 nM of sense 6-toehold 3 gate in presence of 5 nM of Pr-Trigger3 and 15 nM of Pr-STAR6 plasmids. Black lines represent the mean of the replicates, while the shaded regions represent standard deviation of those replicates (n=3). B, D: Average sfGFP/dTomato production rates computed according to the data in the black boxed region from A and C, respectively.

### S6T3 peak production almost matches unregulated plasmid

To further assess the behaviour of S6T3, we compared the traces corresponding to the most efficient concentrations (9.4 nM of STAR/2 nM of trigger) with a positive control that contains only a very strong RBS in its 5’ UTR. As shown in Fig. 9 A, fluorescence intensity between the two constructs shows a similar order of magnitude, even though the plateau phase of S6T3 is reached one hour earlier. We computed the time derivative of the linear phase (Fig. 9 B) and did not observe a significant difference of the sfGFP production rate. This indicates that the translation efficiency of the S6T3 construct is near its optimum. This suggests that the observed earlier plateau is likely an issue of energy limitation related to the transcriptional burden associated with the presence of plasmids encoding the activators’ RNAs. However, the depletion effect is small as both fluorescent values end up on the same order of magnitude. We found this result to be in agreement with previous experiments and model simulations of the influence of non-coding RNA transcription on the translation efficiency in TX-TL (*36*). We then replaced sfGFP by dTomato, a red fluorescent protein, to compare the behaviour of the gate with a different downstream coding sequence. Similar to sfGFP, the order of magnitude of the fluorescence is comparable after 15 hours. However, the production rate of dTomato regulated by S6T3 is about half of the unregulated one. The dTomato sequence greatly varies from sfGFP (43.9% similarity score), and it has been shown that the folding of the mRNA as well as the codon context can influence protein expression (*37*).

Co-expression of protein in TX-TL seems to impose a heavier burden on resources compared to the coexpression of non-coding RNAs. For instance, Shin et al. (*38*) reported a decrease of about 12-fold between the sfGFP production controlled by σ^70^ and an AND-gate based on the enhancer protein Nrtc and σ^54^. Furthermore, STAR and trigger activators are considerably faster than protein activators, which require longer transcription, as well as supplementary steps (translation and maturation). This observation agrees with our expectation to obtain quick logic modules, notably faster than their protein-based counterparts (*39*). We observed here a time delay of the circuit, defined as the time necessary to observe a detectable sfGFP signal, of about 15 minutes. The response time of the P_54_-based AND-gate was found to be around 30 minutes in TX-TL, therefore twice as long as the time required by S6T3. Takahashi et al. (*40*) have found even more striking evidence for faster signal propagation with about 5 minutes per step of a RNA-based cascade in TX-TL against 140 minutes as measured *in vivo* (*41*).

### *In vivo* implementation

Fast prototyping of synthetic circuits *in vitro* often aims to transfer the selected candidates into living organisms. In our study, *coli* TOP10 cells were transformed with two plasmids, one containing the AND-gate controlling the sfGFP-reporter and the other containing one or both activators. Endpoint fluorescent measurements are shown in Fig. 10 A. The nine gates show a very low OFF-state in the absence of either STAR RNA or trigger RNA. In the presence of one RNA activator the sfGFP fluorescence increases only between 3- to 10-fold. If both activating RNAs are present, the reporter fluorescence goes up by more than 50-fold. Comparing the *in vivo* to the *in vitro* measurements, it is possible to see that both measurements are in good agreement with each other (Fig. 10 B). The S6T3 gate has by far the highest ON-state in both the TX-TL and the *E. coli* measurements (614-fold). Those results add a supplementary evidence to the list of demonstrated correlation between cell-free-based and *in vivo* measurements, ranging from the characterisation of single parts (*42*, *43*) to complex dynamic gene circuits (*44*).

**Figure 10:**
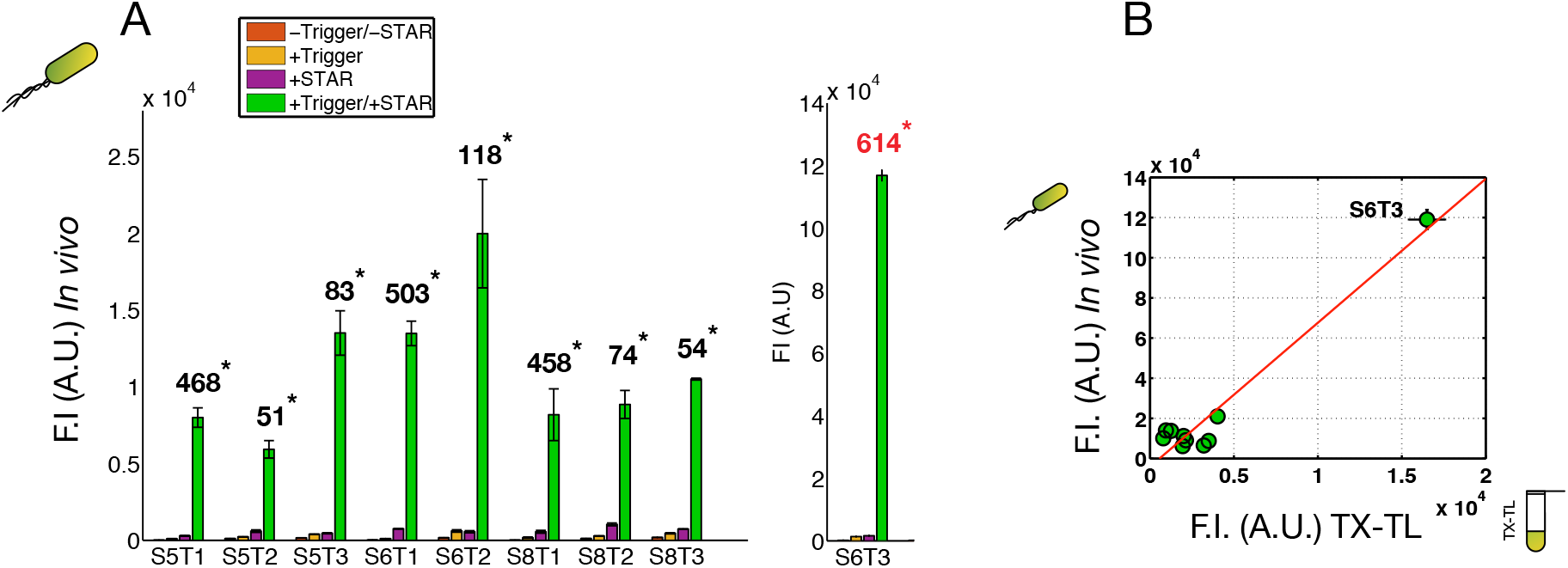
Implementation of the RNA-based gates *in vivo*. (A) Fluorescent measurements of the nine newly combined AND-gates in *E. coli*. The different colors indicate controls without any activators, with only STAR or only trigger. Green bars represent experimental measurements in presence of both STAR and trigger activators. Numbers with * indicate the fold range computed by the ratio between the activated gate and the signal without any activators. Error bars correspond to the standard deviation of three independent measurements. (B) Corresponding relative fluorescence intensities between *in vivo* and *in vitro* for ten RNA-based gates.

## Conclusion

We performed a characterisation of several logic AND-gates built on multi-level RNA activators. To the best of our knowledge, this is the first study which combined the STAR and toehold regulators, two highly efficient RNA activators. We used a TX-TL system together with a robotic liquid handler, which enabled us to perform precise, fast and down-scaled parallel reactions for testing a large range of concentrations. From the resulting dose-response matrices, we determined the saturation concentrations which maximize the dynamic range of our AND-gates. We observed the same saturation behaviour for the gate as for the single parts; the transcriptional activator STAR needed threefold the amount of the translational activator trigger to reach saturation. This agreement between single parts and gates shows the potential of the STAR-toehold system for synthetic circuit design where modularity of parts is desirable.

From the study of our prototype gate, we gained evidence that a combination of Bayesian inference based modeling and TX-TL experiments can lead to a better understanding of synthetic circuit performance. We were able to predict the behaviour of the AND-gate from the parameterisation of its single components. The transcriptional leakage confirmed by the model guided us to use tighter TX-components.

Experimentally, we identified a remarkable AND-gate based on the STAR 6 and toehold 3 parts. Its production rate was as high as for the unregulated sfGFP, showing a great potential for metabolic engineering. We suspect that local co-folding of the RNA regulators hinders the translational process of the remaining gates. In future work, it may be worthwhile to perform more advanced secondary structure analysis and combine it with our ODE population model to obtain better quantitative insights.

In addition, we suggest that our RNA-based gate would be a precious tool to study resource competition in TX-TL. Our device enables the simple, adjustable and independent control of TX- and TL-processes, and is compatible with mRNA monitoring technology (*33*). More work needs to be done in this direction to assess the balance of effects between resource limitation and stoichiometric reagent mismatches (*45*).

We successfully transferred all gates from *in vitro* to *in vivo*, demonstrating the versatility of TX-TL systems for the fast prototyping of RNA-based circuits. Additional studies to transfer the dose-response matrices from the TX-TL systems to an *in vivo* counterpart may help engineer more balanced and efficient circuits in living cells.

Finally, TX-TL based diagnostic platforms with toehold switches have been recently used for the rapid and low-cost detection of specific RNA genome sequences (*46*, *47*). As the STAR activators can be computationally designed (*48*), coupling the toehold sensors with a STAR module could increase the flexibility and sensitivity of the current bio-molecular diagnostic tools.

## Material and Methods

### Cell-free reactions

Cell-free reactions from *E. coli* were purchased (My TX-TL company, 507096) where cell extract and premix containing amino acids were already pre-assembled at a total ratio of 0.75.

### Preparation of cell-free reactions

Before each use, the reactions were thawed at 4 °C and RNase inhibitor (MOLO) was added to the mix (0.01). 1.5 μl of the master mix was dispensed into a clear-bottom 1536-well plate (Greiner) with the help of an automatic dispenser (Eppendorf Repeater E3). The volume was then brought up with plasmid DNA constructs containing the genetic circuits and nuclease-free water (0.5) by using an I-Dot One SC (Dispendix). An Excel sheet computing the DNA concentration range needed for an experiment enables to preview the required amount of total volume needed for each component of the circuit. The microplate was then quickly centrifuged and covered with a transparent sealer (Ampliseal, Greiner). Finally, the microplate was vortexed and centrifuged again before being ready for either kinetic or endpoint measurements.

### Incubation and kinetic measurements

Clear-bottom 1536-well or 384-well black plates containing assembled cell-free reactions and DNA were measured with a PHERAstar FSX (BMG Biotech). The temperature was set to 29 °C for the duration of the experiment (10 hours). The GFP signal was measured every 2 minutes with the direct optic bottom module FI 485/520 (Ex/Em), adjusting gain to 120 with 20 flashes per well and z-height optimised according to each experiment.

### RNA synthesis

For TX-TL measurements of the S5T2 gate with pre-expressed RNAs activators, STAR 5 and trigger 2 were transcribed from PCR-amplified templates, all containing three 5-terminal guanosyl residues to facilitate transcription in vitro using T7 RNA polymerase. For this, two oligonucleotides were designed with an overlap of 35 bp containing the T7 promoter sequence and amplified using Q5 High-Fidelity DNA polymerase (NEB) according to the suppliers instructions. After ethanol precipitation, the DNA template was used for *in vitro* transcription with T7 RNA polymerase (NEB) as reported by the manufacturer. The RNA was gel purified and molarity was determined by spectrophotometric measurements using NanoDrop 1000 Spectrophotometer (Thermo Scientific).

### Endpoint measurements

Clear-bottom 1536-well or 384-well black plates containing assembled cell-free reactions and DNA were placed in an incubator at 29 °C for 10 h. The GFP signal was then measured with the direct optic bottom module FI 485/520 (Ex/Em), adjusting gain to 120 with 20 flashes per well and z-height optimised according to each experiment.

### Assembly of expression constructs

The construction of the plasmid library is based on plasmids previously published: pJBL2801 was a gift from Julius Lucks (Addgene plasmid 71207) and was used for cloning the STARs and triggers; pJBL2807 was a gift from Julius Lucks (Addgene plasmid 71203) and was taken every two minut were e second sum stands for used for cloning the sense, toehold and the nine combined gates. Golden gate and Gibson assembly were used for cloning the inserts with length longer than 100 bps. Q5 site directed mutagenesis kit from NEB was used for cloning the inserts inferiors at 100 bps. Oligos used for the various assemblies or mutagenesis were ordered from Sigma Aldrich. Sanger sequencing verified all constructed plasmids.

Plasmids were prepared using a NucleoBond Extra Midi Plus (Macherey-Nagel) and followed by isopropanol precipitation and eluted with 10 mM Tris at pH=8.5 and quantified using a NanoDrop 1000. Those having an insufficient purity were subject to a Phenol-chloroform extraction followed by a novel step of isopropanol precipitation.

The sequence of each part used is available in the supplemental information (Table S1).

### Flow cytometer analysis

The plasmid containing the regulatory element and the plasmid containing the regulatory RNA were transferred into *E. coli* TOP10 cells by the calcium chloride method. The cells were grown on LB agar plates containing the antibiotics ampicillin and chloramphenicol over night at 37 °C. A single colony was used to inoculate 4 ml of LB containing the needed antibiotics for selection. The cells were grown over night (37 °C, 150 rpm). For sfGFP analysis the cells were diluted 1:10 with PBS into a 96-well plate (flat bottom, Greiner). Fluorescence was analysed by using the CytoFlex S cytometer from Beckmann. For sfGFP excitation a 488 nm laser was used and emission was detected by a 510/20 nm filter. For each sample 25 000 events were measured. Each experiment was repeated three times.

### Parallel tempering

In Bayesian parameter inference, we are concerned with finding the posterior distribution of the parameters ***θ*** included in our model, given the experimental data ***Y***;

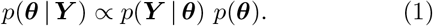

To identify this distribution, we applied MCMC parallel tempering (*49*). This method ensures the exploration of the whole parameter space as well as faster convergence. In the following, we state the main formulas of the algorithm used in this work.

Our log-likelihood is determined by the common assumption of independent normally distributed measurement deviations and we denote it similar to (*26*) by

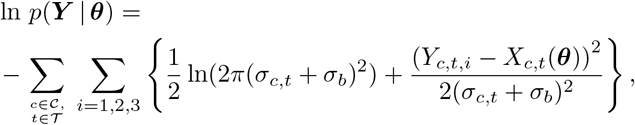

where 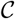 contains the experimental conditions (in our case the different input concentrations of the activator plasmids), and 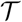 contains the different time points of the measurements (in our case, measurements the three replicates measured for each concentration. The simulated model data of the reporter flourescence intensity are indicated by *X*_*c,t*_(***θ***) due to their dependence on the activator input concentration, the evaluation time, and the parameter ***θ***. The experimental data as well are indicated by the activator concentration, the time point of the measurement and the enumeration of the replicate. From the three replicates for each concentration and each time point, the empirical standard deviation *σ*_*c,t*_ was computed. Further, we set an additional “base noise” *σ*_*b*_ to capture especially the experimental irregularities up to 25 minutes.

As a prior distribution, we choose the uniform distribution on the interval of *±* five orders of magnitude around parameter literature values (e.g. taken from (*50*)).

Samples are drawn from the tempered posteriors, i.e. from the posterior distributions of different temperatures given as follows

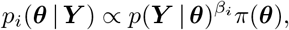

where *β*_*i*_ is the *i*th inverse temperature (thermodynamic parameter). We choose the inverse temperatures according to the following – *geometric* – temperature schedule: Denoting the predefined number of different temperatures by *N*_*t*_, e.g. 50, one further chooses a minimal inverse temperature, here 10^−6^, and identifies *q* ∈ (0, 1) such that 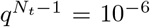. There with the set of inverse temperatures is given by

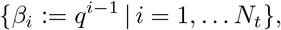

where *β*_1_ = 1 corresponds to the “original temperature”, e.g. to the true posterior distribution.

The acceptance probability for the *i*th chain to switch to a proposed sample ***θ***′ is determined by the well-known uences of the synthetic gates, supplementary data for Metropolis criterion and given by

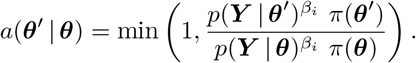

The proposal distribution for a new parameter set is given by the normal distribution with mean equal to the current parameter values and predefined proper variances that are scaled with some predefined constant greater than 1 which scales exponentially with the enumeration of the temperatures, i.e., the highest temperature possesses the largest variance.

Finally, in each sampling step, there is a predefined number of possible swaps of current samples belonging to the differently tempered chains: picking randomly one of the chains, its corresponding current sample can possibly be changed with the current sample from the chain with next higher temperature according to the following swap acceptance probability (*51*, *52*)

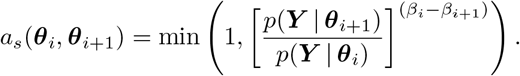

Having characterised the parameter posterior distribution, one can further compute the posterior predictive distribution for a new data point *y′* at time *t′* regarding to an input concentration *c′* via

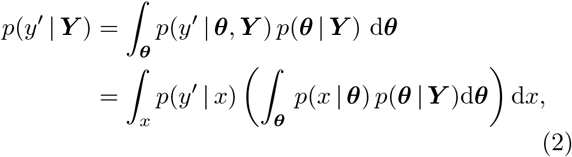

where in our case of a deterministic model, we have *p*(*x* | ***θ***) = *δ*(*x−X*_*c*′, *t*′_ (***θ***)).

## Supporting information

Supplementary informations

## Acknowledgement

Many thanks to Felix Reinhardt for helpful discussions and his C code of the ODE solver to improve performance of our MATLAB sampling framework, to Leo Bronstein for providing us the framework of the parallel tempering implementation, to Lukas Köhs for his input about RNA structures, and to Johannes Kabisch and Niels Schlichting for their help with handling the i-dot robotic platform. This work was supported by the Landesoffensive für wissenschaftliche Exzellenz (LOEWE; initiative to increase research excellence in the state of Hessen, Germany) as part of the LOEWE Schwerpunkt CompuGene.

## Supporting Information Available

Plasmids, sequences of the synthetic gates, supplementary data for experimental and modeling experiments. This material is available free of charge via the Internet at http://pubs.acs.org/.

