## Supplementary informations for "Cell-free prototyping of AND-logic gates based on heterogeneous RNA activators"

### Supplementary information

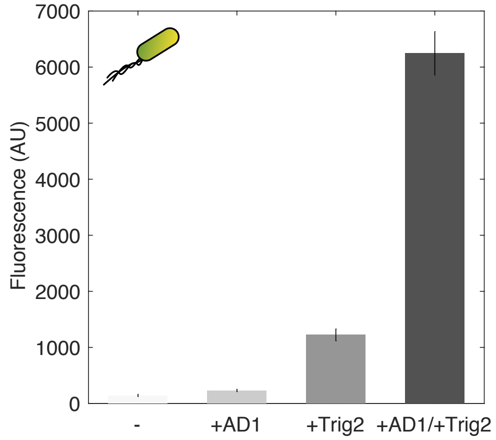


Figure 1: *In vivo* characterisation of the initial gate based on AD1 and toehold 2. Fluorescence endpoint measurements of the gate in top10 *E. coli* cells in presence (+) or absence (-) of the trigger/STAR DNA-encoded activators.

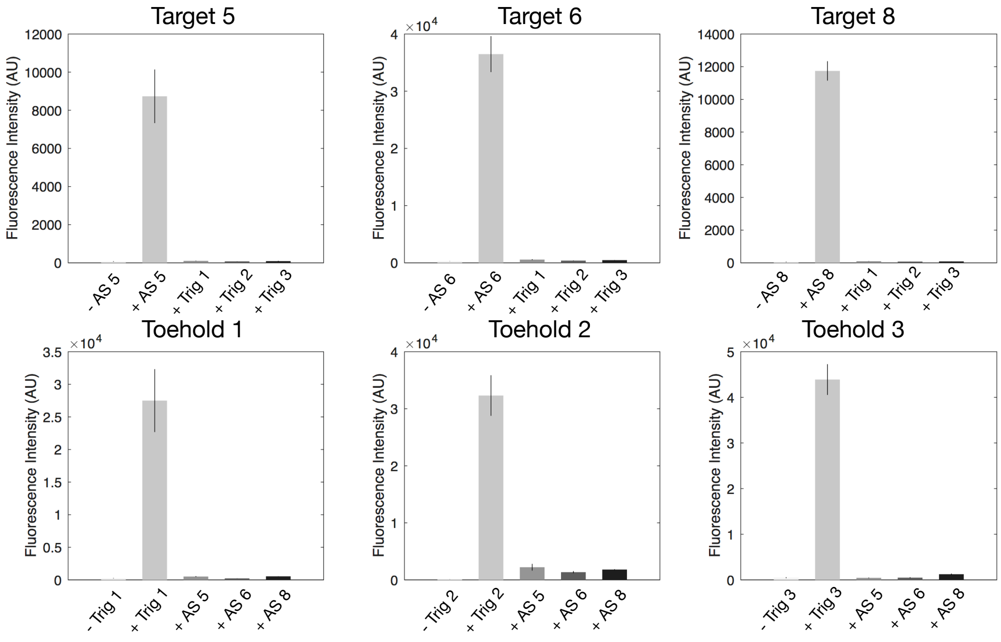


Figure 2: *In vivo* measurements of endpoint fluorescence for each single part, either toehold $(1, 2, 3)$ or STAR target $(5, 6, 8)$, in presence (+) or absence (-) of their cognate or orthogonal trigger or STAR antisense. Error bars correspond to the standard deviation of three independent measurements.

#### Mathematical Modeling of TX- and TL-regulated components

We modeled the transcriptional regulation system STAR AD1-target and the translational regulation system trigger-toehold 2 with parameterized ordinary differential equations (ODE). Our goal was to model the dynamics of the systems with a basic set of equations.

The TX-regulated system was modeled by the following basic reactions; the transcription and degradation of the small mRNA; the binding and unbinding of the small mRNA to the reporter mRNA; the transcription and degradation of the reporter mRNA; the translation of the reporter; and the maturation of the reporter. This resulted in the following system of ODEs

$$\frac{\mathrm{d}R_S}{\mathrm{d}t} = k_S \cdot D_S - d_S \cdot R_S,$$

$$\frac{\mathrm{d}R}{\mathrm{d}t} = k_R \cdot \left[ \frac{R_S^{N_S}}{(R_S^{N_S} + K_S)} + p_S \right] \cdot D_G - d_R \cdot R,$$

$$\frac{\mathrm{d}G}{\mathrm{d}t} = k_{TL} \cdot R - \alpha \cdot G,$$

$$\frac{\mathrm{d}G_m}{\mathrm{d}t} = \alpha \cdot G,$$

with $R_S$ the small mRNA and $D_S$ the corresponding DNA template; $R$ the reporter mRNA with corresponding DNA template $D_G$; $G$ the immature and $G_m$ the mature GFP. Besides the common kinetic parameters like the transcription, degradation, translation, and maturation rates $(k_S, k_R, d_S, d_R, k_{TL}, \alpha)$, the prefactor of the reporter transcription

$$\left[ \frac{R_S^{N_S}}{(R_S^{N_S} + K_S)} + p_S \right]$$

contains a Hill coefficient, $N_S$, the dissociation constant of target-STAR binding, $K_S$, and an *activation probability*, $p_S$. This reflects the TX-regulation property of the system: If the small mRNA is bound to the accruing reporter mRNA, transcsription will be finished. Otherwise, there is still some activation probability for successful transcription caused by incomplete formation of the target terminator.

The ODE system for the TL-regulated system was modeled similarly by the following equations

$$\frac{\mathrm{d}R_T}{\mathrm{d}t} = k_T \cdot D_T - d_T \cdot R_T,$$

$$\frac{\mathrm{d}R}{\mathrm{d}t} = k_R \cdot D_G - d_R \cdot R,$$

$$\frac{\mathrm{d}G}{\mathrm{d}t} = k_{TL} \cdot \left[ \frac{R_T^{N_T}}{(R_T^{N_T} + K_T)} + p_T \right] \cdot R - \alpha \cdot G,$$

$$\frac{\mathrm{d}G_m}{\mathrm{d}t} = \alpha \cdot G.$$

Both parameterized models were then calibrated to the kinetic experimental data via the MCMC method parallel tempering, described in the main text (c.f. Sec. 3 Material and Methods), performing a Bayesian parameter inference. Within this procedure we set $k_S = k_R$ and $k_T = k_R$ in our model, since the transcription rates within both systems should roughly be the same as the same promoters were used.

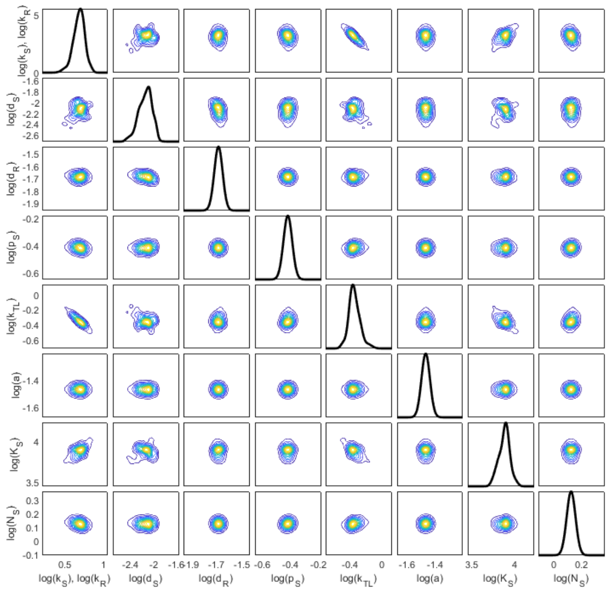


Figure 3: Parameter posterior distributions of the STAR AD1-target system; univariate distributions are on the diagonal, bivariate distributions are on the off-diagonal. The joint posterior was generated with 150 000 samples in the logarithmic space. Trace plots of the parameters were considered to determine the burn-in phase and to ensure that convergence was reached.

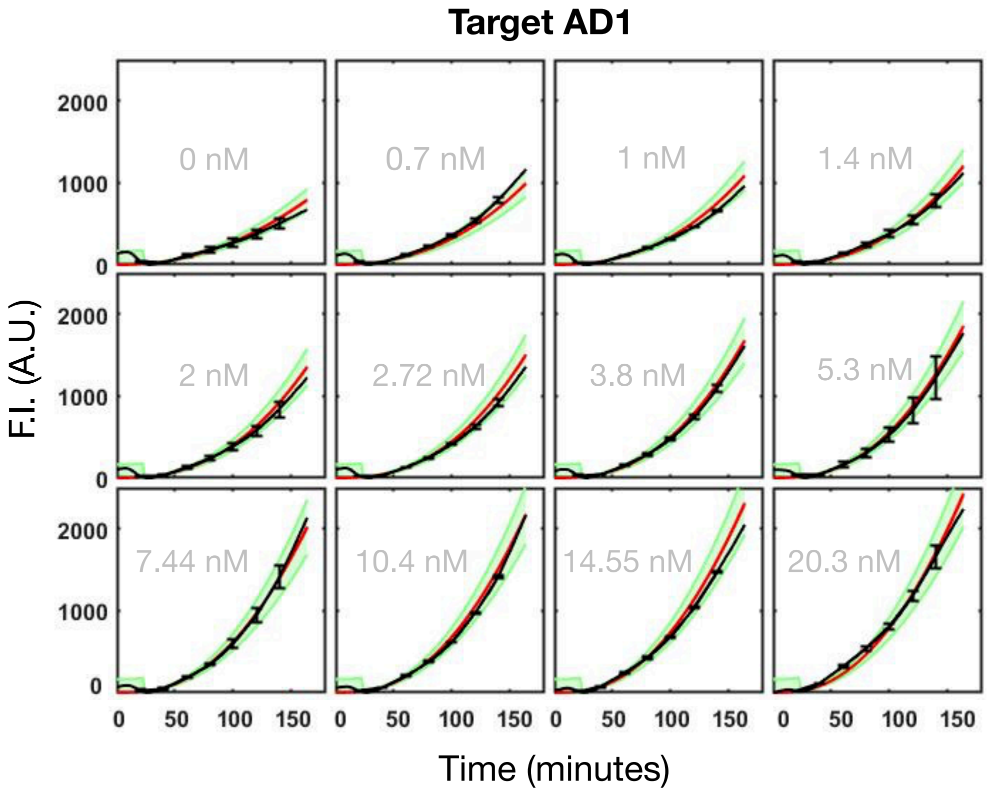


Figure 4: Calibration of single part: posterior predictive distribution of thr STAR AD1-target system, for 2 nM input concentration of target plasmid. Gray in the background: input concentrations of the AD1 plasmid; black: experimental data together with error bars at selected time points; red: median of parameterised model according to identified parameter posterior; green: observation model corresponding to experimentally measured standard deviation and assumption of normally distributed observation errors, 95%- and 5%-quantiles are shown. To capture the experimental irregularities up to 25 minutes, we set our "base noise" parameter higher there.

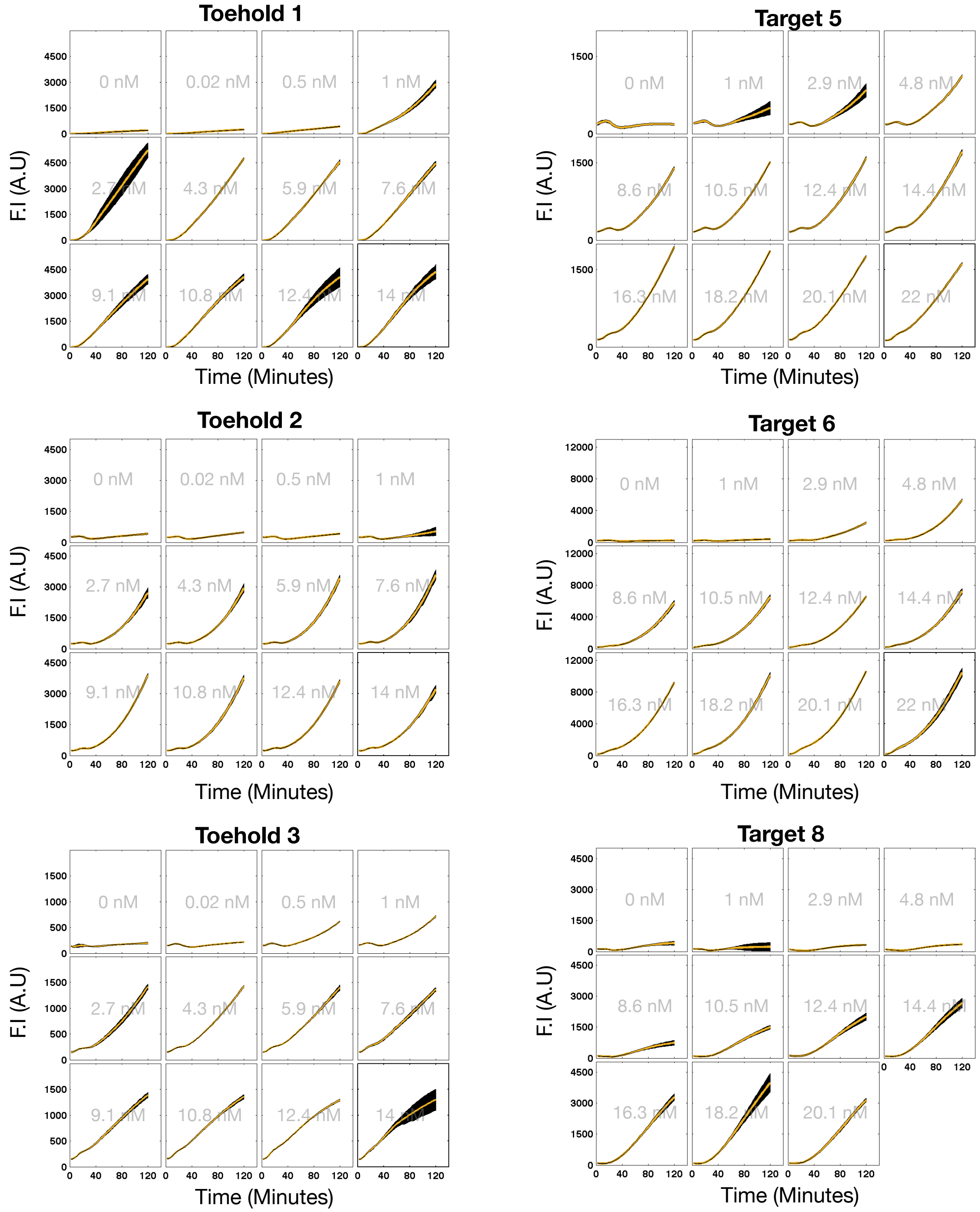


Figure 5: Kinetic characterisation of each individual part in presence of 2nM of reporter template, either toehold or target. Yellow lines correspond to the averaged experimentally measured fluorescence. Black shaded regions represent the experimentally measured standard deviation. Concentrations written in gray in the background correspond to the activator plasmid concentration, either trigger or STAR DNA plasmid.

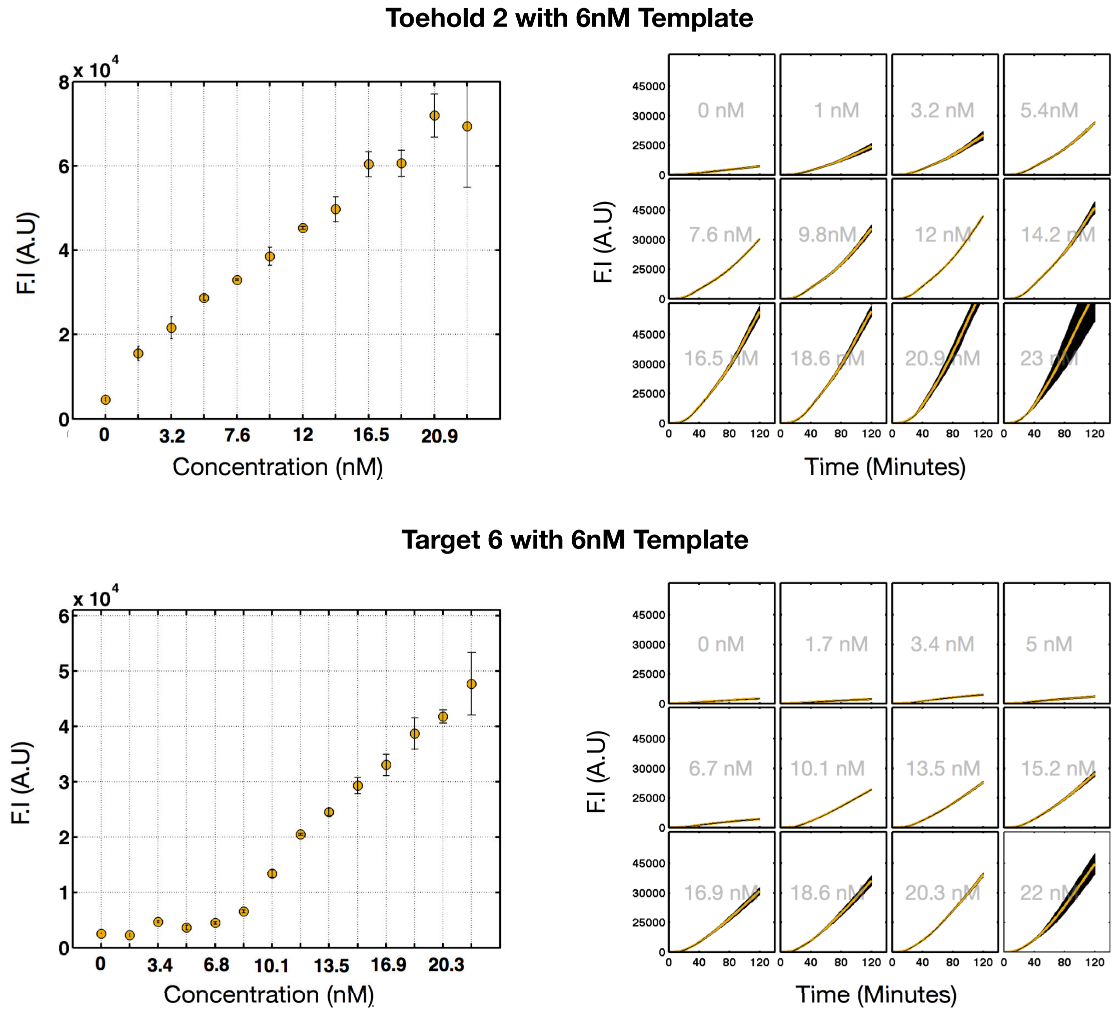


Figure 6: Kinetic characterisation of toehold 2 and target 6 parts in presence of 6nM of target or toehold template. Yellow lines correspond to the averaged experimentally measured fluorescence. Black shaded regions represent the experimentally measured standard deviation. Concentrations written in gray in the background correspond to the activator plasmid concentration, either trigger or STAR encoding plasmid.

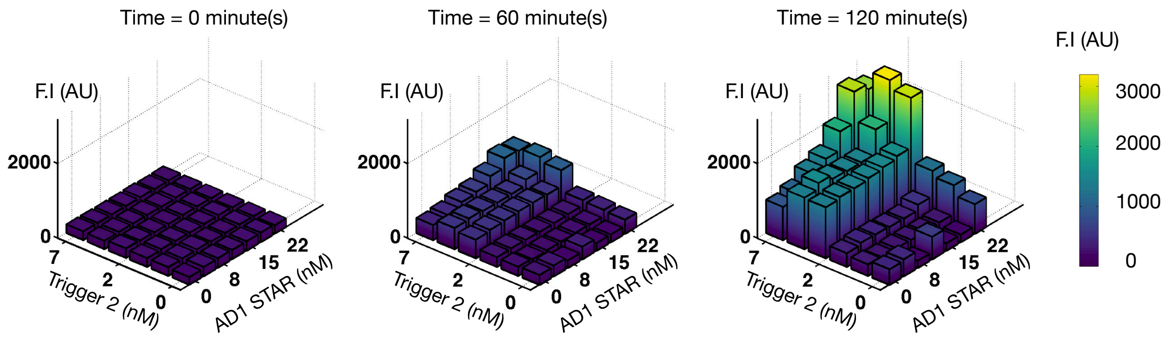


Figure 7: Dose-response fluorescence measurements for the prototyping gate AD1-toehold 2 performed every 60 minutes.

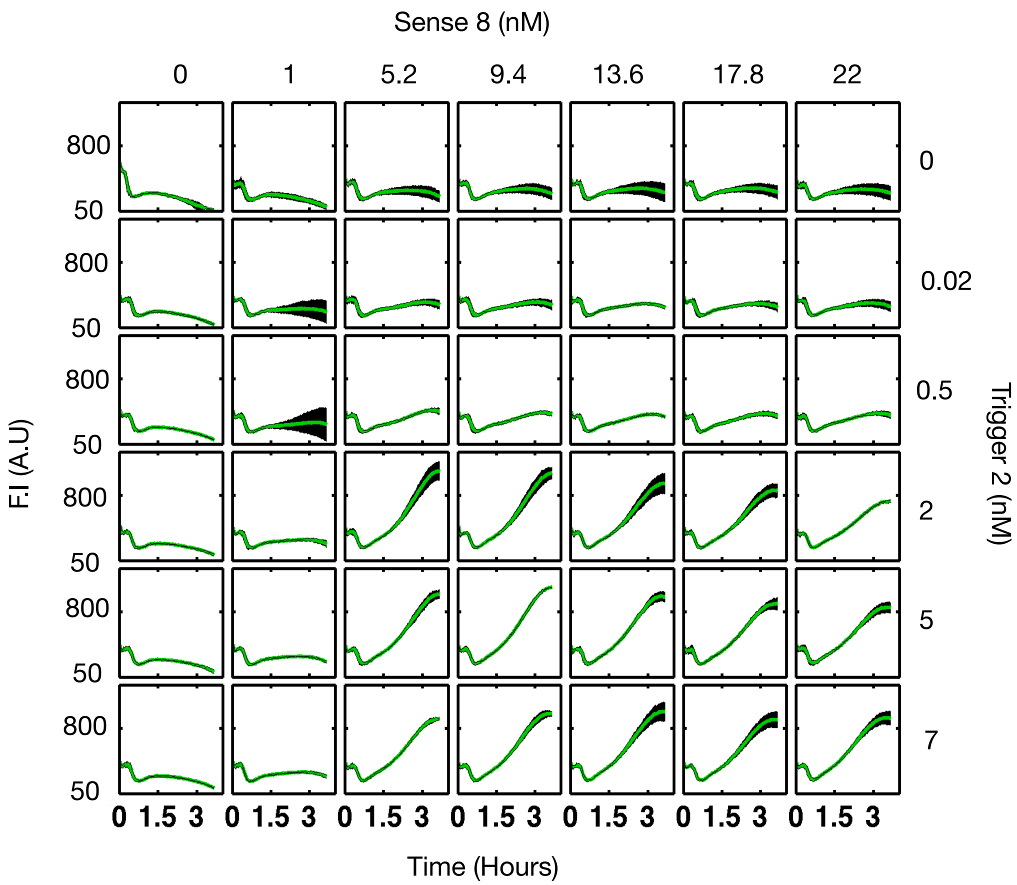


Figure 8: Concentration matrix of TX-TL time course reactions for the sense8-toehold2-SGFP gate (S8T2). Trigger and STAR encoding plasmids were simultaneously titrated from 0 to 7nM and 0 to 22nM, respectively. Green lines correspond to the averaged experimentally measured fluorescence. Black shaded regions represent the experimentally measured standard deviation over three independent measurements.

Table 1: Plasmids used.

| Plasmid Name | Plasmid architecture | Source |
|---|---|---|
| pJBL2801 | J23119 – Target AD1.S5 – RBS – sfGFP – TrrnB – CmR p15A origin | Insert Reference Lucks 2015 Nature Chem Biol |
| pJBL4970 | J23119 – Target5 – RBS – sfGFP – TrrnB – CmR p15A origin | Insert Reference Lucks 2017 nature com |
| pJBL5816 | J23119 – Target6 – RBS – sfGFP – TrrnB – CmR p15A origin | Insert Reference Lucks 2017 nature com |
| pJBL5808 | J23119 – Target8 – RBS – sfGFP – TrrnB – CmR p15A origin | Insert Reference Lucks 2017 nature com |
| pJBL2807 | J23119 – STAR AD1.A5 – T500 – AmpR – ColE1 origin | Insert Reference Lucks 2015 Nature Chem Biol |
| pJBL4971 | J23119 – STAR5 – T500 – AmpR – ColE1 origin | Insert Reference Lucks 2017 nature com |
| pJBL5817 | J23119 – STAR6 – T500 – AmpR – ColE1 origin | Insert Reference Lucks 2017 nature com |
| pJBL5809 | J23119 – STAR8 – T500 – AmpR – ColE1 origin | Insert Reference Lucks 2017 nature com |
| pJBL-wt | J23119 — RBS – sfGFP – TrrnB – CmR p15A origin | this study |
| pJBL-Toehold1 | J23119 – Toehold1 – RBS – sfGFP – TrrnB – CmR p15A origin | this study |
| pJBL-Toehold2 | J23119 – Toehold2 – RBS – sfGFP – TrrnB – CmR p15A origin | this study |
| pJBL-Toehold3 | J23119 – Toehold3 – RBS – sfGFP – TrrnB – CmR p15A origin | this study |
| pJBL-trigger1 | J23119 – trigger1 – T500 – AmpR – ColE1 origin | this study |
| pJBL-trigger2 | J23119 – trigger2 – T500 – AmpR – ColE1 origin | this study |
| pJBL-trigger3 | J23119 – trigger3 – T500 – AmpR – ColE1 origin | this study |
| pJBL-Target AD1.S5-Toehold2 | J23119 – Target AD1.S5 – Toehold2 – sfGFP – TrrnB – CmR p15A origin | this study |
| pJBL-Target5-Toehold1 | J23119 – Target5 – Toehold1 – sfGFP – TrrnB – CmR p15A origin | this study |
| pJBL-Target5-Toehold2 | J23119 – Target5 – Toehold2 – sfGFP – TrrnB – CmR p15A origin | this study |
| pJBL-Target5-Toehold3 | J23119 – Target5 – Toehold3 – sfGFP – TrrnB – CmR p15A origin | this study |
| pJBL-Target6-Toehold1 | J23119 – Target6 – Toehold1 – sfGFP – TrrnB – CmR p15A origin | this study |
| pJBL-Target6-Toehold2 | J23119 – Target6 – Toehold2 – sfGFP – TrrnB – CmR p15A origin | this study |
| pJBL-Target6-Toehold3 | J23119 – Target6 – Toehold3 – sfGFP – TrrnB – CmR p15A origin | this study |
| pJBL-Target8-Toehold1 | J23119 – Target8 – Toehold1 – sfGFP – TrrnB – CmR p15A origin | this study |
| pJBL-Target8-Toehold2 | J23119 – Target8 – Toehold2 – sfGFP – TrrnB – CmR p15A origin | this study |
| pJBL-Target8-Toehold3 | J23119 – Target8 – Toehold3 – sfGFP – TrrnB – CmR p15A origin | this study |
| pJBL-STAR AD1.A5-trigger2 | t500 – trigger2 – J23119 – spacer – J23119 – STAR AD1.A5 – T500 – AmpR – ColE1 origin | this study |
| pJBL-STAR5-trigger1 | t500 – trigger1 – J23119 – spacer – J23119 – STAR5 – T500 – AmpR – ColE1 origin | this study |
| pJBL-STAR5-trigger2 | t500 – trigger2 – J23119 – spacer – J23119 – STAR5 – T500 – AmpR – ColE1 origin | this study |

| Plasmid Name | Plasmid architecture | Source |
|---|---|---|
| pJBL-STAR5-trigger3 | t500 – trigger3 – J23119 – spacer – J23119 – STAR5 – T500 – AmpR – ColE1 origin | this study |
| pJBL-STAR6-trigger1 | t500 – trigger1 – J23119 – spacer – J23119 – STAR6 – T500 – AmpR – ColE1 origin | this study |
| pJBL-STAR6-trigger2 | t500 – trigger2 – J23119 – spacer – J23119 – STAR6 – T500 – AmpR – ColE1 origin | this study |
| pJBL-STAR6-trigger3 | t500 – trigger3 – J23119 – spacer – J23119 – STAR6 – T500 – AmpR – ColE1 origin | this study |
| pJBL-STAR8-trigger1 | t500 – trigger1 – J23119 – spacer – J23119 – STAR8 – T500 – AmpR – ColE1 origin | this study |
| pJBL-STAR8-trigger2 | t500 – trigger2 – J23119 – spacer – J23119 – STAR8 – T500 – AmpR – ColE1 origin | this study |
| pJBL-STAR8-trigger3 | t500 – trigger3 – J23119 – spacer – J23119 – STAR8 – T500 – AmpR – ColE1 origin | this study |
| pJBL-wt(red) | J23119 — RBS – dTomato – TrrnB – CmR p15A origin | this study |
| pJBL-Target6-Toehold3 | J23119 – Target6 – Toehold3 – dTomato – TrrnB – CmR p15A origin | this study |

Table 2: Example Plasmids.

| Name | Sequence [5'->3'] |
|---|---|
| pJBL-STAR5-trigger1 t500, trigger, J23119, STAR, ColE1, AmpR | GAATTCaaaaaaaagcccgcctttcggcgggctttgAGATCCctatcttatcttatctatctcgtttatccctgctttactgact<br>attgcacagaatagtcagtcccactagtattatacctaggactgagctagctgtcaaAGATCTTTAACGGGGTCAT<br>CACGGCTCATCATGCGCCAAACAAATGTGTGCAATACACGCTCGGATGACTGCA<br>TGATGACCGCACTGACTGGGGACAGCAGATCCACCTAAGCCTGTGAGAGAAGCA<br>GACACCCGACAGATCAAGGCAGTTAAATTAAAGATCTttgacagctagctcagtcctaggtat<br>aatactagttgaactgtatacattccccgcaggataggaattgaagatgaaacgatgagacttgggacgaGGATCT<br>caaagcccgccgaaaggcgggcttttttttGGATCCTTACTCGAGTCTAGACTGCAGGCTTCCT<br>CGCTCACTGACTCGCTGCGCTCGGTCGTTCGGCTGCGGCGAGCGGTATCAGCTC<br>ACTCAAAGGCGGTAATACGGTTATCCACAGAATCAGGGGATAACGCAGGAAAGA<br>ACATGTGAGCAAAAGGCCAGCAAAAGGCCAGGAACCGTAAAAAggccgcgttgctggcg<br>tttttccacaggctccgcccccctgacgagcatcacaaaaatcgacgctcaagtcagaggtggcgaaacccgacaggactataaa<br>gataccaggcgtttccccctggaagctccctcgtgcgctctcctgttccgaccctgccgcttaccggatacctgtccgcctttct<br>cccttcgggaagcgtggcgctttctcatagctcacgctgtaggtatctcagttcggtgtaggtcgttcgctccaagctgggctgt<br>gtgcacgaaccccccgttcagcccgaccgctgcgccttatccggtaactatcgtcttgagtccaacccggtaagacacgacttat<br>cgccactggcagcagccactggtaacaggattagcagagcgaggtatgtaggcggtgctacagagttcttgaagtggtggcctaa<br>ctacggctacactagaagaacagtatttggtatctgcgctctgctgaagccagttaccttcggaaaaagagttggtagctcttga<br>tccggcaaacaaaccaccgctggtagcggtggtttttttgtttgcaagcagcagattacgcgcagaaaaaaaggatctcaagaag<br>atcctttgatcttttctacggggtctgacgctcagtggaacgaaaactcacgttaagggattttggtcatgaGATTATCA<br>AAAAGGATCTTCACCTAGATCCTTTTAAATTAAAAATGAAGTTTTAAATCATCT<br>AAAGTATATATGAGTAAACTTGGTCTGACAGTTAccaatgcttaatcagtgaggcacctatctcagc<br>gatctgtctatttcgttcatccatagttgcctgactccccgtcgtgtagataactacgatacgggagggcttaccatctggcccc<br>agtgctgcaatgataccgcgagacccacgctcaccggctccagatttatcagcaataaaccagccagccggaagggccgagcgca<br>gaagtggtcctgcaactttatccgcctccatccagtctattaattgttgccgggaagctagagtaagtagttcgccagttaatag |
| pJBL-STAR5-trigger1 t500, trigger, J23119, STAR, ColE1, AmpR | tttgcgcaacgttgttgccattgctacaggcatcgtggtgtcacgctcgtcgtttggtatggcttcattcagctccggttcccaa<br>cgatcaaggcgagttacatgatcccccatgttgtgcaaaaaagcggttagctccttcggtcctccgatcgttgtcagaagtaagt<br>tggccgcagtgttatcactcatggttatggcagcactgcataattctcttactgtcatgccatccgtaagatgcttttctgtgac<br>tggtgagtactcaaccaagtcattctgagaatagtgtatgcggcgaccgagttgctcttgcccggcgtcaatacgggataatacc<br>gcgccacatagcagaactttaaaagtgctcatCATTGGAAAACGTTCTTCGGGGCGAAAACTCTCA<br>AGGATCTTACCGCTGTTGAGATCCAGTTCGATGTAACCCACTCGTGCACCCAACTG<br>ATCTTCAGCATCTTTTACTTTCACCAGCGTTTCTGGGTGAGCAAAAACAGGAAGGC<br>AAAATGCCGCAAAAAAGGGAATAAGGGCGACACGGAAATGTTGAATACTCATACTC<br>TTCCTTTTTCAATATTATTGAAGCATTTATCAGGGTTATTGTCTCATGAGCGGATA<br>CATATTTGAATGTATTTAGAAAAATAAACAAATAGGGGTTCCGCGCACATTTCCCC<br>GAAAAGTGCCACCTGACGTCTAAGAAACCATTATTATCATGACATTAACCTATAAA<br>AATAGGCGTATCACGAGGCAGAATTTCAGATAAAAAAAATCCTTAGCTTTCGCTAA<br>GGATGATTTCTG |

| Name | Sequence [5'->3'] |
|---|---|
| pJBL-Target5-Toehold1 J23119, Target, toehold, sfGFP, TrrnB, CmR, p15A | GAATTCTAAAGATCTttgacagctagctcagtcctaggtataatactagttcgtcccaagtctcatcgtttcatct<br>tcaattcctatcctgcggggaatgtatacagttcatgtatatattccccgcttttttttGGATCTgggtcttatcttatc<br>tatctcgtttatccctgcatacagaaacagaggagatatgcaatgataaacgagaacctggcggcagcgcaaaag<br>atgagcaaaggagaagaacttttcactggagttgtcccaattcttgttgaattagatggtgatgttaatgggcacaaattttctgtc<br>cgtggagagggtgaaggtgatgctacaaacggaaaactcacccttaaatttatttgcactactggaaaactacctgttccgtgg<br>ccaacacttgtcactactctgacctatggtgttcaatgcttttcccgttatccggatcacatgaaacggcatgactttttcaag<br>agtgccatgcccgaaggttatgtacaggaacgcactatatctttcaaagatgacgggacctacaagacgcgtgctgaagtcaag<br>tttgaaggtgatacccttgttaatcgtatcgagttaaagggtattgattttaaagaagatggaaacattcttggacacaaactc<br>gagtacaactttaactcacacaatgtatacatcacggcagacaaacaaaagaatggaatcaaagctaacttcaaaattcgccac<br>aacgttgaagatggttccgttcaactagcagaccattatcaacaaaatactccaattggcgatggccctgtccttttaccagac<br>aaccattacctgtcgacacaatctgtcctttcgaaagatcccaacgaaaagcgtgaccacatggtccttcttgagtttgtaact<br>gctgctgggattacacatggcatggatgagctctacaaaTAAGCGGCCGCGGATCTgaagcttgggcccgaaca<br>aaaactcatctcagaagaggatctgaatagcgccgtcgaccatcatcatcatcatcattgagtttaaacggtctccagcttggc<br>tgttttggcggatgagagaagattttcagcctgatacagattaaatcagaacgcagaagcggtctgataaaacagaatttgcct<br>ggcggcagtagcgcggtggtcccacctgaccccatgccgaactcagaagtgaaacgccgtagcgccgatggtagtgtggggtct<br>ccccatgcgagagtagggaactgccaggcatcaaataaaacgaaaggctcagtcgaaagactgggcctttcgttttatctgttg<br>tttgtcggtgaactGGATCCTTACTCGAGTCTAGACTGCAGTTGATCGggcacgtaagaggttc<br>caactttcaccataatgaaataagatcactaccgggcgtattttttgagttatcgagattttcaggagctaaggaagctaaaat<br>ggagaaaaaaatcactggatataccaccgttgatatatcccaatggcatcgtaaagaacattttgaggcatttcagtcagttgc<br>tcaatgtacctataaccagaccgttcagctggatattacggcctttttaaagaccgtaaagaaaaataagcacaagttttatcc<br>ggcctttattcacattcttgcccgcctgatgaatgctcatccggaatttcgtatggcaatgaaagacggtgagctggtgatatg<br>ggatagtgttcacccttgttacaccgttttccatgagcaaactgaaacgttttcatcgctctggagtgaataccacgacgattt<br>ccggcagtttctacacatatattcgcaagatgtggcgtgttacggtgaaaacctggcctatttccctaaagggtttattgagaa<br>tatgtttttcgtctcagccaatccctgggtgagtttcaccagttttgatttaaacgtggccaatatggacaacttcttcgcccc<br>cgttttcaccatgggcaaatattatacgcaaggcgacaaggtgctgatgccgctggcgattcaggttcatcatgccgtttgtga<br>tggcttccatgtcggcagaatgcttaatgaattacaacagtactgcgatgagtggcagggcggggcgtaatttgatatcgagct<br>cgcttggactcctgttgatagatccagtaatgacctcagaactccatctggatttgttcagaacgctcggttgccgccgggcgt<br>tttttattGGTGAGAATCCAAGCCTCCGATCAACGTCTCATTTTCGCCAAAAGTTG |
| pJBL-Target5-Toehold1 J23119, Target, toehold, sfGFP, TrrnB, CmR, p15A | GCCCAGGGCTTCCCGGTATCAACAGGGACACCAGGATTTATTTATTCTGCGAAGT<br>GATCTTCCGTCACAGGTATTTATTCGGCGCAAAGTGCGTCGGGTGATGCTGCCAA<br>CTTACTGATTTAGTGTATGATGGTGTTTTTGAGGTGCTCCAGTGGCTTCTGTTTC<br>TATCAGCTGTCCCTCCTGTTCAGCTACTGACGGGGTGGTGCGTAACGGCAAAAGC<br>ACCGCCGGACATCAgcgctagcggagtgtatactggcttactatgttggcactgatgagggtgtcagtgaagtgcttc<br>atgtggcaggagaaaaaaggctgcaccggtgcgtcagcagaatatgtgatacaggatatattccgcttcctcgctcactgactc<br>gctacgctcggtcgttcgactgcggcgagcggaaatggcttacgaacggggcggagatttcctggaagatgccaggaagatact<br>taacagggaagtgagagggccgcggcaaagccgtttttccataggctccgcccccctgacaagcatcacgaaatctgacgctca<br>aatcagtggtggcgaaacccgacaggactataaagataccaggcgtttccccctggcggctccctcgtgcgctctcctgttcct<br>gcctttcggtttaccggtgtcattccgctgttatggccgcgtttgtctcattccacgcctgacactcagttccgggtaggcagt<br>tcgctccaagctggactgtatgcacgaaccccccgttcagtccgaccgctgcgccttatccggtaactatcgtcttgagtccaa<br>cccggaaagacatgcaaaagcaccactggcagcagccactggtaattgatttagaggagttagtcttgaagtcatgcgccggtt<br>aaggctaaactgaaaggacaagttttggtgactgcgctcctccaagccagttacctcggttcaaagagttggtagctcagagaa<br>ccttcgaaaaaccgccctgcaaggcggttttttcgttttcagagcaagagattacgcgcagaccaaaacgatctcaagaagatc<br>atcttattaatcagataaaatatttCTAGATTTCAGTGCAATTTATCTCTTCAAATGTAGCACC<br>TGAAGTCAGCCCCATACGATATAAGTTGTAATTCTCATGTTTGACAGCTTATCAT<br>CGATAAGCTTCCGATGGCGCGCCGAGAGGCTTTACACTTTATGCTTCCGGCT |

Table 3: Part sequences

| Name | Sequence [5'->3'] |
|---|---|
| Promoter J23119 | TTGACAGCTAGCTCAGTCCTAGGTATAATACTAGT |
| Terminator t500 | CAAAGCCCGCCGAAAGGCGGGCTTTTTTTT |
| Terminator TrrnB | GAAGCTTGGGCCCGAACAAAAACTCATCTCAGAAGAGGATCTGAATAGCGC<br>CGTCGACCATCATCATCATCATCATTGAGTTTAAACGGTCTCCAGCTTGGC<br>TGTTTTGGCGGATGAGAGAAGATTTTCAGCCTGATACAGATTAAATCAGAA<br>CCAGAAGCGGTCTGATAAAACAGAATTTGCCTGGCGGCAGTAGCGCGGTGG<br>TCCCACCTGACCCCATGCCGAACTCAGAAGTGAAACGCCGTAGCGCCGATG<br>GTAGTGTGGGGTCTCCCCATGCGAGAGTAGGGAACTGCCAGGCATCAAATA<br>AAACGAAAGGCTCAGTCGAAAGACTGGGCCTTTCGTTTTATCTGTTGTTTG TCGGTGAACT |

| Name | Sequence [5’->3’] |
|---|---|
| AmpR | CCAATGCTTAATCAGTGAGGCACCTATCTCAGCGATCTGTCTATTTCGTTC<br>ATCCATAGTTGCCTGACTCCCCGTCGTGTAGATAACTACGATACGGGAGGG<br>CTTACCATCTGGCCCCAGTGCTGCAATGATACCGCGAGACCCACGCTCACC<br>GGCTCCAGATTTATCAGCAATAAACCAGCCAGCCGGAAGGGCCGAGCGCAG<br>AAGTGGTCCTGCAACTTTATCCGCCTCCATCCAGTCTATTAATTGTTGCCG<br>GGAAGCTAGAGTAAGTAGTTCGCCAGTTAATAGTTTGCGCAACGTTGTTGC<br>CATTGCTACAGGCATCGTGGTGTCACGCTCGTCGTTTGGTATGGCTTCATT<br>CAGCTCCGGTTCCCAACGATCAAGGCGAGTTACATGATCCCCCATGTTGTG<br>CAAAAAAGCGGTTAGCTCCTTCGGTCCTCCGATCGTTGTCAGAAGTAAGTT<br>GGCCGCAGTGTTATCACTCATGGTTATGGCAGCACTGCATAATTCTCTTAC<br>TGTCATGCCATCCGTAAGATGCTTTTCTGTGACTGGTGAGTACTCAACCAA<br>GTCATTCTGAGAATAGTGTATGCGGCGACCGAGTTGCTCTTGCCCGGCGTC<br>AATACGGGATAATACCGCGCCACATAGCAGAACTTTAAAAGTGCTCAT |
| CmR | GGCACGTAAGAGGTTCCAACTTTCACCATAATGAAATAAGATCACTACCG<br>GGCGTATTTTTTGAGTTATCGAGATTTTCAGGAGCTAAGGAAGCTAAAAT<br>GGAGAAAAAAATCACTGGATATACCACCGTTGATATATCCCAATGGCATC<br>GTAAAGAACATTTTGAGGCATTTCAGTCAGTTGCTCAATGTACCTATAAC<br>CAGACCGTTCAGCTGGATATTACGGCCTTTTTAAAGACCGTAAAGAAAAA<br>TAAGCACAAGTTTTATCCGGCCTTTATTCACATTCTTGCCCGCCTGATGA<br>ATGCTCATCCGGAATTTCGTATGGCAATGAAAGACGGTGAGCTGGTGATA<br>TGGGATAGTGTTCACCCTTGTTACACCGTTTTCCATGAGCAAACTGAAAC<br>GTTTTCATCGCTCTGGAGTGAATACCACGACGATTTCCGGCAGTTTCTAC<br>ACATATATTCGCAAGATGTGGCGTGTTACGGTGAAAACCTGGCCTATTTC<br>CCTAAAGGGTTTATTGAGAATATGTTTTTCGTCTCAGCCAATCCCTGGGT<br>GAGTTTCACCAGTTTTGATTTAAACGTGGCCAATATGGACAACTTCTTCG<br>CCCCCGTTTTCACCATGGGCAAATATTATACGCAAGGCGACAAGGTGCTG<br>ATGCCGCTGGCGATTCAGGTTCATCATGCCGTTTGTGATGGCTTCCATGT<br>CGGCAGAATGCTTAATGAATTACAACAGTACTGCGATGAGTGGCAGGGCG<br>GGGCGTAATTTGATATCGAGCTCGCTTGGACTCCTGTTGATAGATCCAGT<br>AATGACCTCAGAACTCCATCTGGATTTGTTCAGAACGCTCGGTTGCCGCC<br>GGGCGTTTTTTATT |
| ColE1 origin | GGCCGCGTTGCTGGCGTTTTTCCACAGGCTCCGCCCCCCTGACGAGCATC<br>ACAAAAATCGACGCTCAAGTCAGAGGTGGCGAAACCCGACAGGACTATAA<br>AGATACCAGGCGTTTCCCCCTGGAAGCTCCCTCGTGCGCTCTCCTGTTCC<br>GACCCTGCCGCTTACCGGATACCTGTCCGCCTTTCTCCCTTCGGGAAGCG<br>TGGCGCTTTCTCATAGCTCACGCTGTAGGTATCTCAGTTCGGTGTAGGTC<br>GTTCGCTCCAAGCTGGGCTGTGTGCACGAACCCCCCGTTCAGCCCGACCG<br>CTGCGCCTTATCCGGTAACTATCGTCTTGAGTCCAACCCGGTAAGACACG<br>ACTTATCGCCACTGGCAGCAGCCACTGGTAACAGGATTAGCAGAGCGAGG<br>TATGTAGGCGGTGCTACAGAGTTCTTGAAGTGGTGGCCTAACTACGGCTA<br>CACTAGAAGAACAGTATTTGGTATCTGCGCTCTGCTGAAGCCAGTTACCT<br>TCGGAAAAAGAGTTGGTAGCTCTTGATCCGGCAAACAAACCACCGCTGGT<br>AGCGGTGGTTTTTTTGTTTGCAAGCAGCAGATTACGCGCAGAAAAAAAGG<br>ATCTCAAGAAGATCCTTTGATCTTTTCTACGGGGTCTGACGCTCAGTGGA<br>ACGAAAACTCACGTTAAGGGATTTTGGTCATGA |
| p15A origin | GCGCTAGCGGAGTGTATACTGGCTTACTATGTTGGCACTGATGAGGGTGT<br>CAGTGAAGTGCTTCATGTGGCAGGAGAAAAAAGGCTGCACCGGTGCGTCA<br>GCAGAATATGTGATACAGGATATATTCCGCTTCCTCGCTCACTGACTCGC<br>TACGCTCGGTCGTTCGACTGCGGCGAGCGGAAATGGCTTACGAACGGGGC<br>GGAGATTTCCTGGAAGATGCCAGGAAGATACTTAACAGGGAAGTGAGAGG<br>GCCGCGGCAAAGCCGTTTTTCCATAGGCTCCGCCCCCCTGACAAGCATCA<br>CGAAATCTGACGCTCAAATCAGTGGTGGCGAAACCCGACAGGACTATAAA<br>GATACCAGGCGTTTCCCCCTGGCGGCTCCCTCGTGCGCTCTCCTGTTCCT<br>GCCTTTCGGTTTACCGGTGTCATTCCGCTGTTATGGCCGCGTTTGTCTCA<br>TTCCACGCCTGACACTCAGTTCCGGGTAGGCAGTTCGCTCCAAGCTGGAC<br>GTATGCACGAACCCCCCGTTCAGTCCGACCGCTGCGCCTTATCCGGTAAC<br>TATCGTCTTGAGTCCAACCCGGAAAGACATGCAAAAGCACCACTGGCAGC<br>AGCCACTGGTAATTGATTTAGAGGAGTTAGTCTTGAAGTCATGCGCCGGT<br>TAAGGCTAAACTGAAAGGACAAGTTTTGGTGACTGCGCTCCTCCAAGCCA<br>GTTACCTCGGTTCAAAGAGTTGGTAGCTCAGAGAACCTTCGAAAAACCGC<br>CCTGCAAGGCGGTTTTTTCGTTTTCAGAGCAAGAGATTACGCGCAGACCA<br>AAACGATCTCAAGAAGATCATCTTATTAATCAGATAAAATATTT |

| Name | Sequence [5'->3'] |
|---|---|
| sfGFP | ATGAGCAAAGGAGAAGAACTTTTCACTGGAGTTGTCCCAATTCTTGTTGA ATTAGATGGTGATGTTAATGGGCACAAATTTTCTGTCCGTGGAGAGGGTG AAGGTGATGCTACAAACGGAAAACTCACCCTTAAATTTATTTGCACTACT GGAAAACTACCTGTTCCGTGGCCAACACTTGTCACTACTCTGACCTATGG TGTTCAATGCTTTTCCCGTTATCCGGATCACATGAAACGGCATGACTTTT TCAAGAGTGCCATGCCCGAAGGTTATGTACAGGAACGCACTATATCTTTC AAAGATGACGGGACCTACAAGACGCGTGCTGAAGTCAAGTTTGAAGGTGA TACCCTTGTTAATGTATCGAGTTAAAGGGTATTGATTTTAAAGAAGATGG AAACATTCTTGGACACAAACTCGAGTACAACTTTAACTCACACAATGTAT ACATCACGGCAGACAAACAAAAGAATGGAATCAAAGCTAACTTCAAAATT CGCCACAACGTTGAAGATGGTTCCGTTCAACTAGCAGACCATTATCAACA AAATACTCCAATTGGCGATGGCCCTGTCCTTTTACCAGACAACCATTACC TGTCGACACAATCTGTCCTTTCGAAAGATCCCAACGAAAAGCGTGACCAC ATGGTCCTTCTTGAGTTTGTAACTGCTGCTGGGATTACACATGGCATGGA TGAGCTCTACAAA |
| spacer | AGATCTTTAACGGGGTCATCACGGCTCATCATGCGCCAAACAAATGTGTG CAATACACGCTCGGATGACTGCATGATGACCGCACTGACTGGGGACAGCA GATCCACCTAAGCCTGTGAGAGAAGCAGACACCCGACAGATCAAGGCAGT TAAATTAAAGATCT |
| Target AD1.S5 | AGTTTTTACAGTGAATTGTTTTAATTAGTTGTATAAATGTTGGAGCAGCG GGGAATGTATACAGTTCATGTATATATTCCCCGCTTTTTTTTT |
| Target5 | TCGTCCCAAGTCTCATCGTTTCATCTTCAATTCCTATCCTGCGGGGAATG TATACAGTTCATGTATATATTCCCCGCTTTTTTTTT |
| Target6 | CCAGTCATCAAGTCAGTCCAGTCAAAGTTTCCGTCGTTCAGCGGGGAATGT ATACAGTTCATGTATATATTCCCCGCTTTTTTTTT |
| Target8 | CCATCCTCAATCTCTACCTACTCTCACTTACTCTTATCCTGCGGGGAATGT ATACAGTTCATGTATATATTCCCCGCTTTTTTTTT |
| Toehold1 | GGGTCTTATCTTATCTATCTCGTTTATCCCTGCATACAGAAACAGAGGAGA TATGCAATGATAAACGAGAACCTGGCGGCAGCGCAAAAG |
| Toehold2 | GGGAGTTTGATTACATTGTCGTTTAGTTTAGTGATACATAAACAGAGGAGA TATCACATGACTAAACGAAACCTGGCGGCAGCGCAAAAG |
| Toehold3 | GGGATCTATTACTACTTACCATTGTCTTGCTCTATACAGAAACAGAGGAGA TATAGAATGAGACAATGGAACCTGGCGGCAGCGCAAAAG |
| STAR AD1.A5 | TGAACTGTATACATTCCCCGCTGCTCCAACATTTATACAACTAATTAAAAC AATTCACTGTAAAAACT |
| STAR5 | TGAACTGTATACATTCCCCGCAGGATAGGAATTGAAGATGAAACGATGAGA CTTGGGACGA |
| STAR6 | TGAACTGTATACATTCCCCGCTGAACGACGGAAACTTTGACTGGACTGACT TGATGACTGG |
| STAR8 | TGAACTGTATACATTCCCCGCAGGATAAGAGTAAGTGAGAGTAGGTAGAGA TTGAGGATGG |
| trigger1 | GGGACTGACTATTCTGTGCAATAGTCAGTAAAGCAGGGATAAACGAGATAG ATAAGATAAGATAG |
| trigger2 | GGGACAGATCCACTGAGGCGTGGATCTGTGAACACTAAACTAAACGACAAT GTAATCAAACTAAC |
| trigger3 | GGGTGATGGGACATTCCGATGTCCCATCAATAAGAGCAAGACAATGGTAAG TAGTAATAGATAAG |

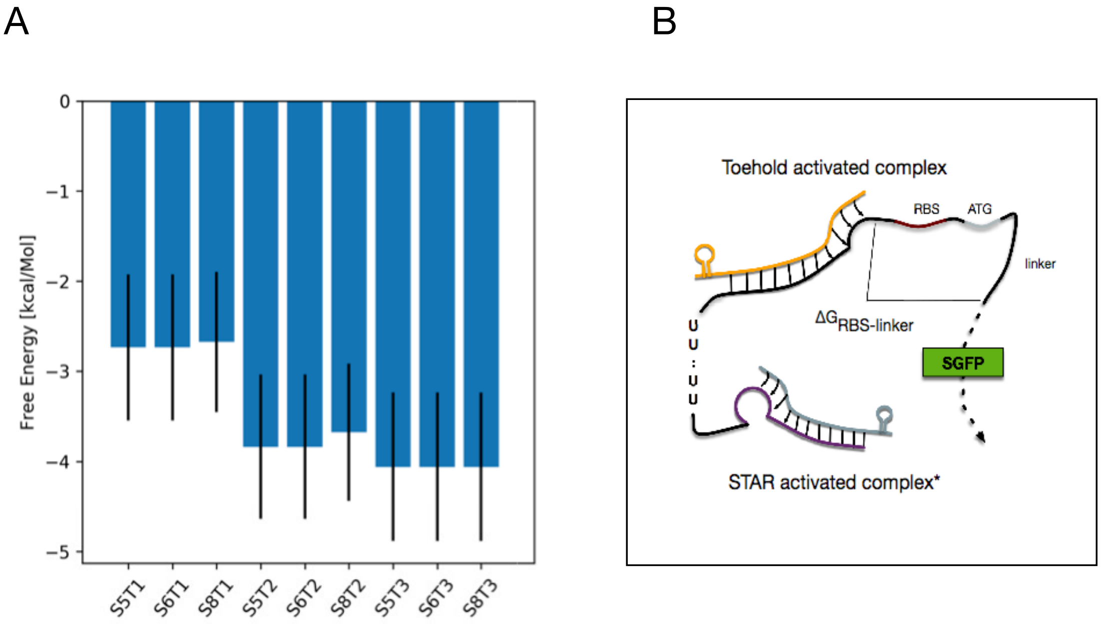


Figure 9: Free Energy of the complexes RBS-linker regions. (A) Mean Free Energies of the RBS-linker regions for a range of suboptimal structures. All suboptimal structures in an energy gap of $\Delta G = 2.5$ kCal/Mol have been computed with NUPACK's `subopt` method. Subsequently, we exclude all structures that have a tail length of $< 47$ nt, i.e. all structures where the toehold-trigger complex covers at least one RBS-nucleotide. The means are calculated over the remaining suboptimal structures. Error bars: one standard deviation. (B) Graphical representation displaying the RBS-linker region for which the Free Energies are shown.

Table 4: Minimum Free Energy (MFE) structures of the tails of all target-toehold constructs in presence of the cognate trigger activator. First lines are structures in dot-parenthesis notation, second lines show the sequences and the third lines the functional elements of interest (`R`: Ribosome Binding Sites; `ATG`: start codons). MFE structures have been computed for the compound consisting of the toehold-trigger and the complex starting from the first nucleotide downstream of the STAR-activated complex using NUPACK's `subopt` method. The tail of a STAR-toehold sequence is here defined as the 3' part of the construct RNA downstream of the last bond to the trigger RNA, corresponding to the "RBS-linker" (see Supp. Fig. 9 B).

| |
|---|
| S8T3<br>`............(((...(((...........)))...))).(((....))).....`<br>`ATACAGAAACAGAGGAGATATAGAATGAGACAATGGAACCTGGCGGCAGCGCAAAAG`<br>`----------RRRRRRRR------ATG------------------------------` |
| S6T3<br>`............(((...(((...........)))...))).(((....))).....`<br>`ATACAGAAACAGAGGAGATATAGAATGAGACAATGGAACCTGGCGGCAGCGCAAAAG`<br>`----------RRRRRRRR------ATG------------------------------` |
| S5T3<br>`............(((...(((...........)))...))).(((....))).....`<br>`ATACAGAAACAGAGGAGATATAGAATGAGACAATGGAACCTGGCGGCAGCGCAAAAG`<br>`----------RRRRRRRR------ATG------------------------------` |
| S8T2<br>`.........((...((....))...))...............(((....))).....`<br>`ATACATAAACAGAGGAGATATCACATGACTAAACGAAACCTGGCGGCAGCGCAAAAG`<br>`----------RRRRRRRR------ATG------------------------------` |
| S6T2<br>`.........((...((....))...))...............(((....))).....`<br>`ATACATAAACAGAGGAGATATCACATGACTAAACGAAACCTGGCGGCAGCGCAAAAG`<br>`----------RRRRRRRR------ATG------------------------------` |
| S5T2<br>`.........((...((....))...))...............(((....))).....`<br>`ATACATAAACAGAGGAGATATCACATGACTAAACGAAACCTGGCGGCAGCGCAAAAG`<br>`----------RRRRRRRR------ATG------------------------------` |
| S8T1<br>`............(((.......................))).(((....))).....`<br>`ATACAGAAACAGAGGAGATATGCAATGATAAACGAGAACCTGGCGGCAGCGCAAAAG`<br>`----------RRRRRRRR------ATG------------------------------` |
| S6T1<br>`............(((.......................))).(((....))).....`<br>`ATACAGAAACAGAGGAGATATGCAATGATAAACGAGAACCTGGCGGCAGCGCAAAAG`<br>`----------RRRRRRRR------ATG------------------------------` |
| S5T1<br>`............(((.......................))).(((....))).....`<br>`ATACAGAAACAGAGGAGATATGCAATGATAAACGAGAACCTGGCGGCAGCGCAAAAG`<br>`----------RRRRRRRR------ATG------------------------------` |
